# Memory CD4 T cell subset organization in the female reproductive tract is regulated via the menstrual cycle through CCR5 signaling

**DOI:** 10.1101/2022.10.01.510445

**Authors:** Alison Swaims-Kohlmeier, Alexander N. Wein, Felica P. Hardnett, Anandi N. Sheth, Zheng-Rong Tiger Li, M. Elliot Williams, Jessica Radzio-Basu, HaoQiang Zheng, Chuong Dinh, Lisa B. Haddad, Elizabeth M.B. Collins, Jenna L. Lobby, Kirsten Kost, Sarah L. Hayward, Igho Ofotokun, Rustom Antia, Christopher D. Scharer, Anice C. Lowen, J. Gerardo Garcia-Lerma, Jacob E. Kohlmeier

**Affiliations:** Department of Gynecology and Obstetrics, Emory University School of Medicine, Atlanta, GA, USA; Department of Microbiology and Immunology, Emory University School of Medicine, Atlanta, GA, USA; Division of HIV/AIDS Prevention, Centers of Disease Control and Prevention, Atlanta, GA, USA; Department of Medicine, Division of Infectious Diseases, Emory University School of Medicine and Grady Health System, Atlanta, GA, USA; Department of Biology, Emory University, Atlanta, GA, USA

**Author notes:** Correspondence: Alison Swaims-Kohlmeier, PhD, or Jacob E. Kohlmeier, PhD, 1510 Clifton Road, RRC 3133, Atlanta, GA, 30322, Email: (A.S-K.), (J.E.K).

## Abstract

Despite their importance for immunity against sexually transmitted infections (STIs), the composition of the female reproductive tract (FRT) memory CD4 T cell population in response to changes in the local tissue environment during the menstrual cycle remains poorly defined. Here we show that across humans, non-human primates (NHP), and mice, FRT CD4 T cells comprise distinct subsets corresponding to migratory memory (T_MM_) and resident memory (T_RM_) cells. T_MM_ display tissue-itinerant trafficking characteristics, restricted FRT tissue distribution, with distinct transcriptional properties and effector responses to infection. CD4 T cell subset fluctuations synchronized with cycle-driven proinflammatory changes within the local tissue environment and oral administration of a CCR5 antagonist inhibited cycle phase-specific migratory T cell surveillance. This study provides novel insights into the dynamic nature of FRT memory CD4 T cells and identifies the menstrual cycle as a key regulator of memory T cell defense at the site of STI exposure.

**Summary:** The menstrual cycle regulates memory T cell surveillance.

## Introduction

The World Health Organization (WHO) reported in 2016 that more than 1 million sexually transmitted infections (STIs) are acquired per day (*1*), and in the United States cases of STIs reached a record high in 2019 (*2*). STIs can lead to a range of health complications in women including pelvic inflammatory disease (PID), cervical cancer, and stillbirths, highlighting the need for defining correlates of immune protection in the female reproductive tract (FRT) mucosa. Many studies have shown that tissue-resident memory CD4 T cells (T_RM_) are critical for the rapid elimination of various viral and bacterial pathogens in the FRT (*3–6*). For example, memory CD4 T cells in the FRT are necessary for protection against STIs including *Chlamydia trachomatis* (*C. trachomatis*) and herpes simplex virus type 2 (HSV-2) (*7, 8*). However, CD4 T cell localization in the FRT may also increase susceptibility to human immunodeficiency virus (HIV) infection (*9*). These discordant roles for FRT memory CD4 cells in disease outcomes underscore the importance of identifying the mechanisms that dictate memory CD4 T cell subset localization and function within the FRT microenvironment.

Similar to other mucosal tissues, CD4 T cell migration into the FRT is regulated by specific chemokine receptors and integrins (*10, 11*). In response to an infection, CCR5, CXCR3, and α4β1 have been identified as important for antigen-specific T cell trafficking to both the upper and lower FRT (*12–14*). Whereas in the absence of infection (homeostatic conditions) it is thought that very little T cell trafficking into the FRT occurs, and local recall responses depend upon tissue-anchored resident memory T cells (*15, 16*). However, in contrast to other mucosal tissues, the FRT undergoes extensive and repeated tissue remodeling events as a result of menstruation, and the impact of these processes on the FRT memory CD4 T cell population and cellular immunity against STIs remains poorly defined.

The menstrual cycle has a substantial influence on immune properties throughout the FRT (*9*). In preparation for potential blastocyst implantation following ovulation, the local immune environment becomes more tolerogenic. However, in the absence of fertilization within the luteal phase of the cycle the process of immune-mediated tissue remodeling of the decidualized endometrium is initiated in order to induce menstruation (*17*). Although the relative phases of the menstrual cycle and in particular the fluctuations in sex-hormone production such as progesterone [P4] and estradiol [E2] have been previously linked with increased susceptibility to STIs such as *C. trachomatis* and HIV (*18–21*), the precise immune mechanisms underlying this change in STI-acquisition risk are not well-understood (*22*). Moreover, while the menstrual cycle regulates immunologic changes throughout the FRT our understanding of how such alterations impact pre-existing memory T cells in the tissue and immune protection are limited by the scarcity of model systems which experience menstruation (*23*). T cell populations in the FRT have been reported to exhibit some variation based upon cycle phase (*9, 21*). Importantly, we previously reported on a CCR5+ CD38+ memory CD4 T cell population in the FRT that fluctuated over the cycle(*24*). Notably, this FRT CD4 T cell population was also enriched for expression of CCR7, indicative of migratory memory T cells (T_MM_) with the potential to migrate from the tissue back into circulation (*25, 26*), but how FRT CD4 T cell surveillance is impacted by the menstrual cycle are unclear.

To better understand CD4 T cell immune surveillance in the FRT and the dynamics of memory CD4 T cells in response to menstrual cycle regulation, we investigated memory CD4 T cell subsets in humans, non-human primates (NHP), and mouse models. We found that while T_RM_ show little variation over time, the FRT memory CD4 T cell population becomes dominated by infiltrating T_MM_ during the luteal phase of the menstrual cycle. RNA sequencing and cytokine analysis showed that FRT T_MM_ are a distinct population from both T_RM_ throughout the FRT and lymph node memory T cells. Furthermore, we discovered that T_MM_-specific responses were effective in immune defense against *Chlamydia muridarum* (*C. Muridarum*) infection, contributing to early control of infection with more rapid bacterial clearance. To identify the mechanisms driving T_MM_ infiltration into the FRT we investigated cell trafficking properties and using mixed bone marrow chimeras we identified CCR5 as critical for CD4 T cell recruitment into the lower FRT. Using the pig-tailed macaque (*Macaca nemestrina*) model to longitudinally monitor FRT CD4 T cell subsets over the course of multiple reproductive cycles, we identified periodic T_MM_ immigration events corresponding to fluctuations in both local and systemic CCR5 chemokines, and T_MM_ migration was inhibited by oral administration of the CCR5 antagonist drug, maraviroc. Taken together, these results shed new light on how the menstrual cycle regulates the dynamic organization of the FRT memory CD4 T cell population and cellular immunity against STIs.

## Results

### Distinct FRT CD4 T cell populations identified by migratory properties

Our previous detection of memory CD4 T cell fluctuations occurring over the menstrual cycle suggested that the cycle might influence migratory T cell immune surveillance(*24*). To better investigate T cell trafficking properties at the FRT barrier site, cervicovaginal lavage (CVL) and blood samples from healthy women participants were measured for cellular expression of chemokine receptors and integrins critical for T cell trafficking into various peripheral tissue sites, including the FRT (*27*). As shown (**Fig. 1A-C**), in comparison to participant-matched CD4 T cells from PBMC, we found an increased frequency of CCR5, CCR6, α4β7, and CCR4 expression from memory CD4 T cells in the CVL samples, consistent with previous reports (*24, 28–30*). Typical of T cells residing in nonlymphoid tissue, lymphoid and endothelial adhesion proteins L-Selectin and P-Selectin were less likely to be expressed on FRT memory CD4 T cells as compared with PBMC CD4 T cells. The fractalkine receptor, CX3CR1, which functions in CD8 T cell inflammatory trafficking (*31*) was also increased on FRT memory CD4 T cells compared with PBMC CD4 T cells, though to our knowledge this has not been previously reported.

**Figure 1.**
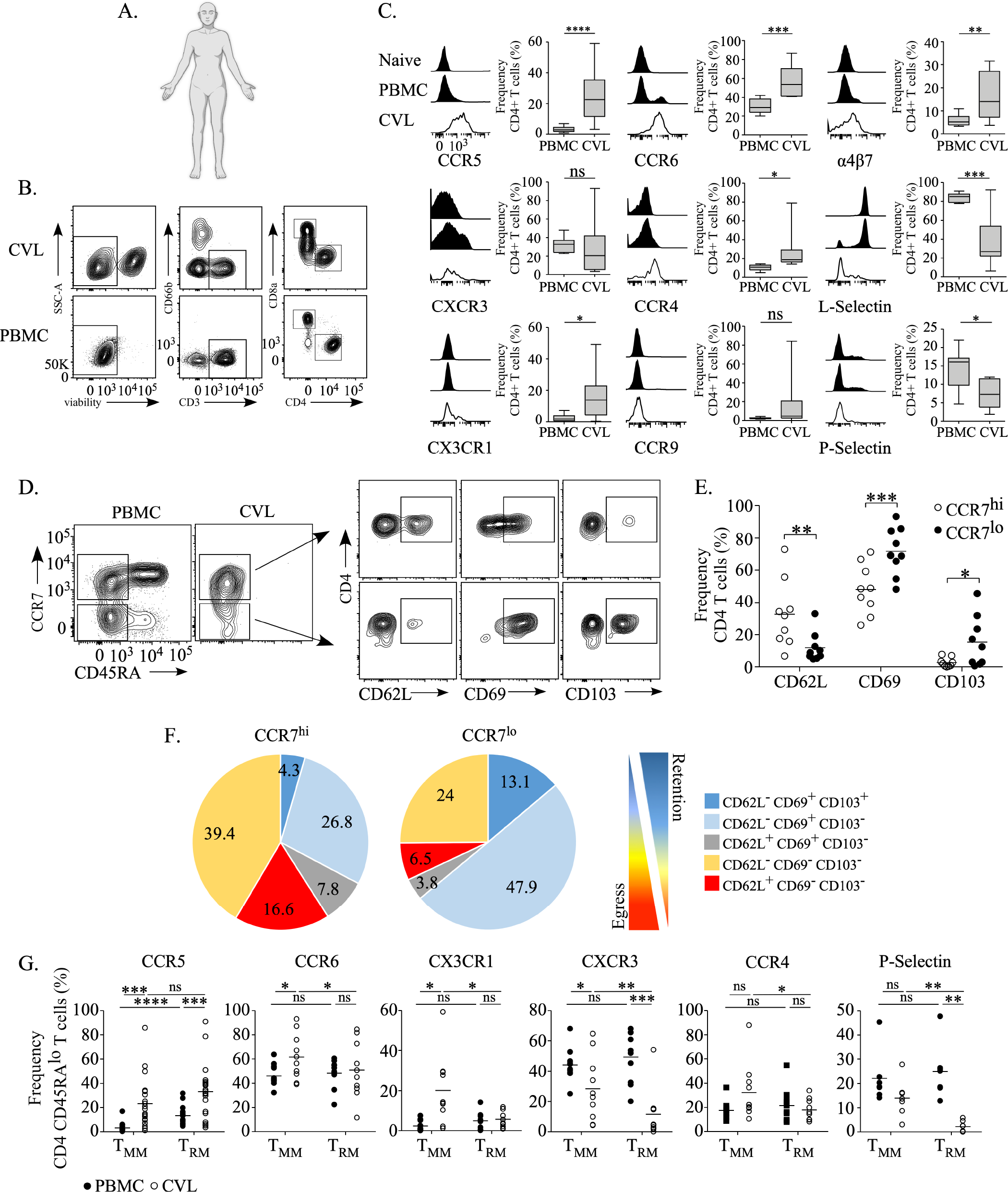
Distinct populations of FRT CD4 T cells identified by expression of tissue residence and trafficking markers. **(A)**. Profile of the biologic female human form created with BioRender.com. **(B).** Representative gating strategy for characterization of CD4 T cell populations enriched from CVL (top panels) compared with matched-PBMC (bottom panels). Viable lymphocytes are gated to distinguish T cells from granulocytes (CD66b) prior to CD4 gating. **(C).** The expression of indicated chemokine receptors or integrins are illustrated by representative histograms (left panels) of naïve gated (CCR7^hi^ CD45RA^lo^) CD4 T cells from PBMC (top histogram). Total PBMC CD4 T cells (center histogram black) and CVL CD4 T cells (bottom histogram white) were compared by the mean frequency of cell surface expression (right panels) of chemokine receptors or integrins (N≥9 per group, graphs depicted as box and whiskers indicating the standard deviation [SD] and mean). **(D).** Representative staining plot illustrating CCR7 and CD45RA gating from PBMC and CVL CD4 T cells followed by expression of CD62L, CD69, and CD103 from either CCR7^hi^ CD45RA^lo^ or CCR7^lo^ CD45RA^lo^ populations of CVL CD4 T cells. **(E).** The frequency of CD62L, CD69, and CD103 expression measured on CCR7^hi^ CD4 T cells (white dots) and CCR7^lo^ CD4 T cells (black dots) from CVL, (N=9, graph depicted as a scatter plot with mean line). **(F).** Boolean pie charts of CD62L, CD69, and CD103 co-expression frequency from either CCR7^hi^ CD45RA^lo^ or CCR7^lo^ CD45RA^lo^ populations of CVL CD4 T cells. **(G).** Chemokine receptors or integrins measured from T_MM_ (CCR7^hi^ CD45RA^lo^) or T_RM_ (CCR7^lo^ CD45RA^lo^) populations from CVL or matched-PBMC CD4 T cells. (C, E, G). (N≥9 per group, graph depicted by scatter plot with mean line). *p≤0.05, **p<0.01, ***p<0.001, ****p<0.0001.

Prior characterization of CD4 T cells from the lower FRT barrier region identified a prominent population consistent with recirculating or migratory memory T cells (T_MM_) (*24*). T_MM_ have been previously identified in human skin (*32*) and mechanistic studies in mice demonstrated that T_MM_ migrate from cutaneous sites into lymph nodes in a CCR7-dependent fashion under steady-state conditions (*25*). Thus, we asked whether we would observe distinct trafficking characteristics by distinguishing FRT CD4 T cells by T_MM_ and T_EM_ (effector memory: CCR7^lo^ CD45RA^lo^). CVL CD4 T cells were gated for CCR7 and CD45RA expression to distinguish T_MM_ or T_EM_ with matched central memory (T_CM_: CCR7^hi^ CD45RA^lo^) or T_EM_ CD4 T cells from PBMC (**Fig. 1D**) and assessed for expression of molecules associated with tissue residence (CD69 and CD103) or egress (CD62L) (**Fig. 1E**). T_MM_ expressed an increased frequency of CD62L with reduced levels of CD69 and CD103 compared with T_EM_. Analysis of FRT CD4 T cell subsets by boolean gating of CD62L, CD69, and CD103 co-expression comparing CCR7^hi^ and CCR7^lo^ FRT memory CD4 T cells showed distinct patterns, with FRT T_MM_ exhibiting a profile more consitent with tissue egress (CD62L^+/-^, CD69^-^, CD103^-^) (*15, 25*) and FRT CD4 T_EM_ populations predominantly expressing classic tissue retention or residence characteristics (CD62L^-^, CD69^+^, CD103^+/-^; FRT T_EM_ from CVL are herein referred to as FRT T_RM_) (*33*) (**Fig. 1F**). As T_MM_ may function in immune surveillance between peripheral and lymphoid tissues, in contrast to T_RM_ which are more stationary in tissues, we applied the trafficking profile to compare FRT T_MM_ and T_RM_ (**Fig. 1G**). T_MM_ cells expressed increased frequencies of CCR6, CX3CR1, CXCR3, CCR4, and P-Selectin in comparison to T_RM_. No differences in the expression of CCR9 and α4α7 were found comparing FRT CD4 T cell subsets (**Fig. S1**). Both FRT T_MM_ and T_RM_ expressed an increased frequency of CCR5 compared with T_CM_ and T_EM_ populations in blood, though there was no difference between FRT CD4 T cell subsets. These results show that T_MM_ predominantly account for trafficking molecule expression observed on CD4 T cells at the lower FRT barrier, though both T_RM_ and T_MM_ are enriched for CCR5 expression.

### Functional distinctions expressed by CD4 T cells based upon anatomic location within the FRT

T cells residing within mucosal barriers can provide rapid and protective responses against infection as compared with circulating memory T cells (*34–37*), but whether FRT CD4 T cells can further exhibit distinct immune functions based upon subset classification or localization within a specific FRT region is unknown. Thus, to investigate the functional response by CD4 T cell subsets, we first assessed cytokine responses to a non-specific stimulus (**Fig. S2**). Both T_MM_ and T_RM_ mostly produced hallmark Th1 cytokines IFNγ and IL-2 in response to immune activation with little to no IL-17 detected. (**Fig. S2B,C**). However, T_MM_ exhibited a higher frequency of IL-2-production with a lower frequency of IFNγ. Notably, both subset populations also predominantly produced the pro-inflammatory cytokine TNFα while not producing any detectable IL-10 (data not shown) indicating that these cells were unlikely to be enriched for regulatory or tolerogenic T cell populations. Next, to investigate FRT CD4 T cell subset responses to antigen-specific stimulation we looked at the effector response to SHIV_162P3_ using NHP tissues (**Fig. S3A**). FRT T cells enriched from CVL at three weekly intervals of sample collection (post peak viremia) were stimulated with overlapping SHIV_162P3_ envelope peptides ex-vivo this was followed by peptide stimulation from FRT T cells enriched from FRT lamina propria at necropsy (**Fig. S3 B-D**). Similar to human samples, we identified a prominent population of T_MM_ present in CVL samples (**Fig. S3C**). Furthermore, we uniquely found high amounts of IL-2 producing CD4 T cells detected from both the CVL and lamina propria compared with little to no IL-2 responses detected from PBMC CD4 T cells (**Fig. S3D**). Similar levels of IFNγ and TNFα production by CD4 T cells were detected across tissues (**Fig. S4**). These data show that FRT T_MM_ are detected in NHP as well as humans and exhibit functional distinctions between FRT T_RM_ and T_EM_ subsets.

Anatomical regions of the FRT (vagina, cervix, and uterus) comprise distinct mucosal barriers based in part upon epithelial composition, mucus production, and tissue-derived signaling interactions (*9*). This suggests that FRT memory CD4 T cells might exhibit different functional characteristics based upon specific tissue localization. Therefore, to better understand memory CD4 T cell distribution throughout the FRT, we measured immune cell populations from pig-tailed macaques (*Macaca nemestrina*) (**Fig. 2A**), a NHP which experience endocrine controlled menstruation over lunar cycles similarly to humans (*38*). To first determine the structural positioning of T_MM_ within the FRT we performed immunofluorescence microscopy from vaginal and cervical tissue sections. From vaginal tissues, we identified T cells (TCRαβ expressing cells) within the basal layer of the vaginal epithelium frequently orienting in aggregates (**Fig. 2B**). Similarly, CCR7+ T cells were identified within the basal layer of vaginal epithelium and progressing towards the apical surface (**Fig. 2C**). CCR7+ T cells were not detected within cervical epithelium (data not shown), suggesting that FRT T_MM_ are preferentially localized to vaginal tissues.

**Figure 2.**
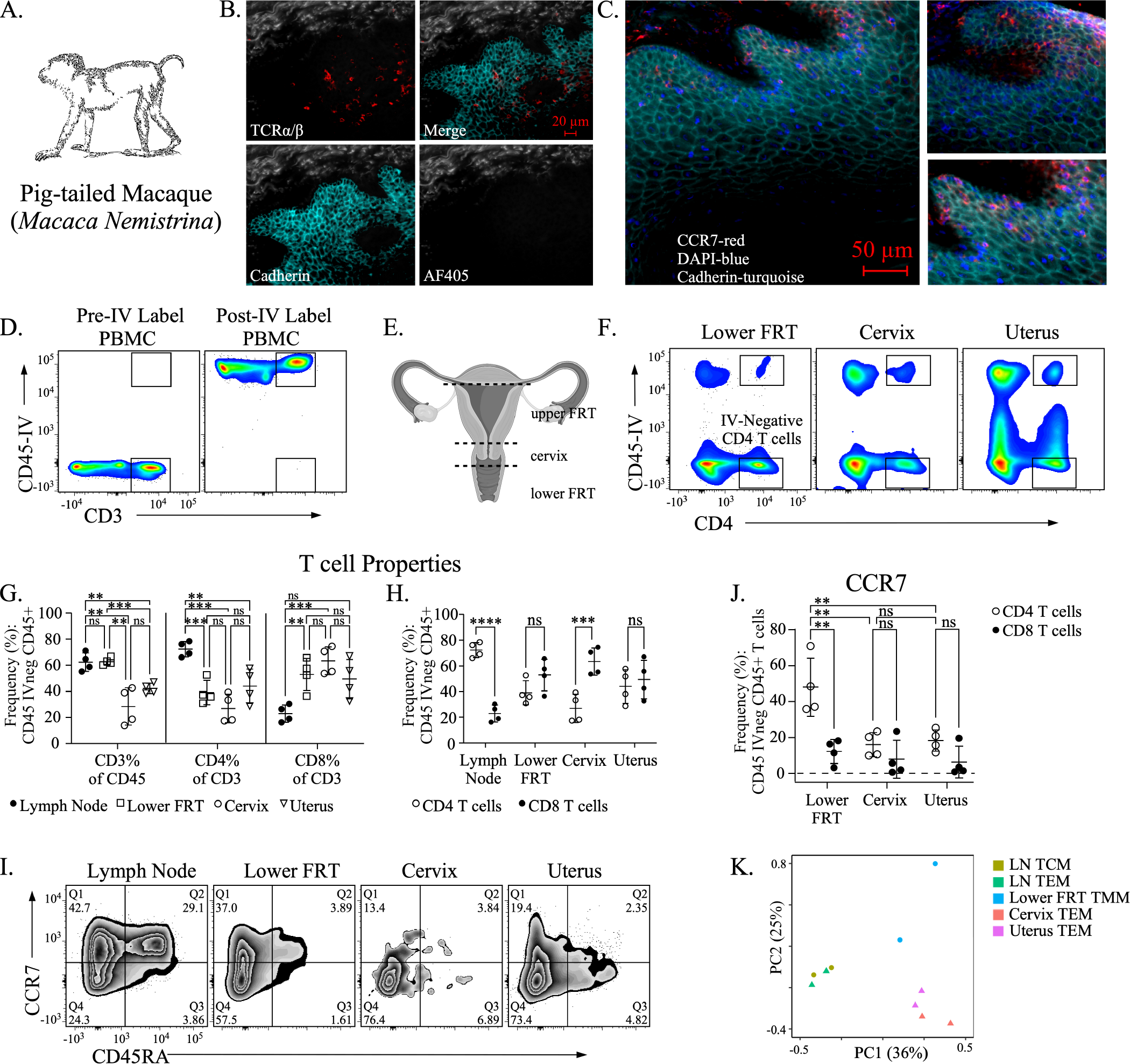
FRT CD4 T cell subset characterization from pig-tailed macaques. **(A).** Cartoon depiction of a pig-tailed macaque. **(B, C).** Fluorescence microscopy image of T cells localized in the lower FRT epithelium of macaques. T cells are identified by TCRα/μ (B) and CCR7 (C) expression while epithelial cells are identified by cadherin expression (B, C). **(D).** Cell flow cytometry plots depicting T cells prior to (left panel) and following IV administration of αCD45 (right panel). **(E).** Cartoon of a macaque or human female reproductive tract with major anatomic regions indicated (created with BioRender.com). **(F).** Representative cell flow plots depicting IV discrimination of CD4 T cells from FRT tissue compartments. **(G, H).** Scatter dot plot graphs with mean line comparing either the T cell populations (G) or the frequency of CD4 and CD8 T cells (H) from the from lymph nodes, the lower FRT, cervix, or uterus. **(I).** Representative flow cytometry cell plots for discriminating memory CD4 T cell subset by expression of CD45RA and CCR7 by tissue. **(J).** Scatter dot plot graphs with mean line comparing CCR7 expression from memory CD4 and CD8 T cells by tissue. (K). PCA plots of mechanically sorted memory CD4 T cells from indicated tissues following RNA-seq. (G, H, J). Means evaluated by multiple comparisons. N=4 *p≤0.05, **p<0.01, ***p<0.001, ****p<0.0001.

Next, to perform T cell characterizations throughout the FRT (including the uterus) that would enable discrimination of true tissue-localized cell populations from contamination by circulating cells within intersecting blood vessels, we performed intravascular (IV) labeling prior to tissue collection **(Fig. 2D-F)** (*39*). Previous immune characterizations from human FRT tissues (performed from biopsy or hysterectomy) were unable to exclude cells from the vasculature or a potential impact of hormonal contraceptive treatment on immune populations prior to tissue removal. Within the lower FRT (vaginal tissues), T cells were the predominant immune cell population among the total (CD45+) leukocyte pool similar to the draining lymph node (dLN) and in contrast to the cervical and uterine regions **(Fig. 2G)**. Furthermore, as compared with T cells from the dLN, all FRT tissues exhibited a lower frequency of CD4 T cells and a higher frequency of CD8 T cells **(Fig. 2H)**. A direct comparison of the CD4 to CD8 T cell ratio by tissue site only showed differences with the dLN and cervix. As expected for the dLN, CD4 T cells comprised the predominant population of the T cell pool while in the cervix, CD8 T cells were the majority. Next, we measured for the FRT CD4 T cell subsets by CCR7 expression **(Fig. 2I)**. By comparing the FRT compartments we identified T_MM_ specifically enriched within the lower FRT **(Fig. 2J)**. Notably, CCR7 expression was mainly restricted to CD4 T cells localized within the lower FRT region, whereas CD8 T cells throughout the FRT (including the lower region) expressed little to no CCR7, consistent with tissue resident T cells (*40*). Together these data show that T_MM_ populations preferentially reside in the vaginal tissues and are concentrated at the vaginal epithelium.

Then, To determined whether the transcriptional program of memory CD4 T cell subsets differed by anatomic region the FRT, we performed bulk RNA sequencing from mechanically sorted IV-negative CD4 T cells including T_MM_ (**Fig. 2K**). A principal component analysis (PCA) showed that memory CD4 T cell populations were transcriptionally distinguishable by tissue region. Specifically, T_MM_ expressed distinct gene signatures as compared with both cervical and uterine memory CD4 T cells as well as T_CM_ and T_EM_ from the dLN, indicating that this population had a unique transcriptional signature.

### Transcriptional profiling of FRT memory CD4 T cell subsets

Despite their functional and phenotypic differences, the transcriptional programs of T_MM_ and T_RM_ in the FRT have not been explored, and the large number of CCR7^hi^ memory CD4 T cells in the FRT raised the possibility that the T_MM_ subset may be comprised of, or related to, T_CM_. Thus, we performed RNA sequencing to compare the transcription profiles of T_MM_ and T_RM_ from the lower FRT with lymph node CD4 T_CM_ from uninfected rhesus macaques (Fig. 3). Differentially expressed genes (DEGs) [false discovery rate (FDR) < 0.05, absolute log_2_fold change (log_2_FC) > 1] indicated that both T_MM_ and T_RM_ subsets had a distinct transcriptional profile compared to T_CM_ cells, and hierarchical clustering separated the subsets into two primary groups; T_CM_ in cluster I, and both T_RM_ and T_MM_ in cluster 2 (**Fig. 3A**). Gene Ontogeny (GO) analysis indicated that cluster II was enriched for expression of genes involved in T-helper type-1 responses, specifically pathways associated with TNFa, IFNy, and IL2 production, in addition to pathways associated with cell migration and T cell differentiation (**Fig 3B, Supplemental Table 1**). Similar to our finding that IFNy and IL2 are differentially produced from T_MM_ and T_RM_ subsets upon stimulation, GSEA analysis found significant enrichment of these pathways in both T_MM_ and T_RM_ subsets compared to T_CM_ (**Fig. 3C, D**). While the overall enrichment of T_MM_ and T_RM_ subsets were similar when compared to T_CM_, differences in expression of specific genes within the IL-2 signaling (**Fig 3E**) and IFNy response (**Fig 3F**) pathways were observed, suggesting some distinct transcriptional differences between T_RM_ and T_MM_ subsets. To further assess the relatedness of CD4 T_MM_ and T_RM_ transcriptomes, we compared expression of genes associated with the core T_RM_ transcriptional signature (*41*). Compared to T_CM_, genes associated with the core T_RM_ signature were differentially expressed in both T_RM_ and T_MM_ subsets (**Fig 3G, H**). However, a direct comparison of upregulated core T_RM_ genes between T_RM_ and T_MM_ subsets showed a significant enrichment of these genes within the T_RM_ subset (**Fig 3I**). Together, these data show that while CD4 T_RM_ and T_MM_ subsets in the FRT have largely overlapping transcriptional signatures, they differ in expression of key genetic programs that define distinct lineages of memory CD4 T cell subsets.

**Figure 3.**
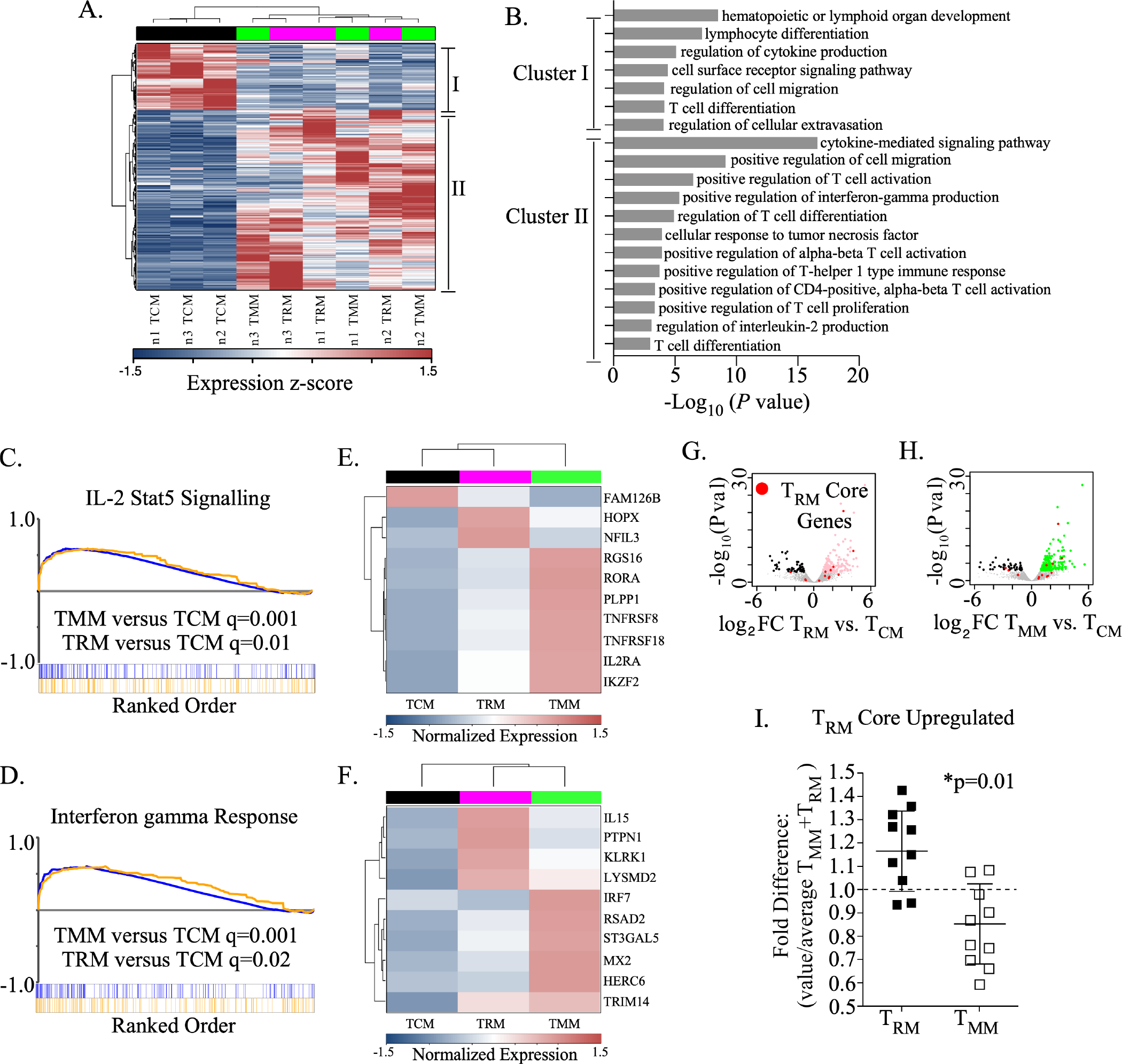
Transcriptional profiling of CD4 T cell subsets in the lower FRT. **(A) .** Heatmap showing z-score normalized expression and hierarchical clustering of 582 DEG. Cluster I and II denote genes upregulated in T_CM_ or T_MM_/T_RM_ cells, respectively. **(B).** Bar plot of gene ontology analysis for pathways enriched in Cluster I or II genes from **A**. GSEA analysis comparing the enrichment of **(C)** IL2-STAT5 Signaling or **(D)** Interferon gamma Response pathways in T_RM_ vs. T_CM_ and T_MM_ vs T_CM_ cells. The FDR q-value of each comparison is noted. Heatmap showing the z-score normalized expression of **(E)** 10 genes from the IL-2 STAT5 Signaling gene set and **(F)** 10 genes from the Interferon gamma Response gene set. The mean expression for each group is shown. Volcano plot showing the log_2_ fold change (FC) in gene expression versus the -log_10_ of the p-value for **(G)** T_RM_ vs. T_CM_ and **(H)** T_MM_ vs T_CM_ cells. The location of core T_RM_ signature genes are noted in red. **(I).** Fold difference in expression levels of upregulated T_RM_ core signature genes comparing sorted CD4 FRT T_RM_ and T_MM_.

### FRT CD4 T_MM_ protect against chlamydia challenge

In order to elucidate the specific contribution of FRT T_MM_ surveillance to immune defense against infection, we utilized laboratory mouse models (**Fig. 4**). First, to identify whether FRT T_MM_ were present in mice we evaluated FRT chemotaxis to CCR7 ligands (as previously reported, commercial mouse-specific CCR7 antibodies did not effectively discriminate CCR7 expression from T cells)(*42*). CD4 T cells enriched from the upper (uterine), cervical (tissue region containing the cervix), or lower (vaginal) region of the FRT (**Fig. 4A,B**) were assessed for CCR7-specific trafficking following CCL21 (a CCR7 chemokine) transwell assay (**Fig. 4C**). To exclude contamination by any circulating T cells, IV labeling was perfomed prior to tissue harvests (*43*). This analysis showed that the lower FRT CD4 T cell populations exhibited the greatest chemotactic index to CCL21. Some migration was detected from cervical-enriched CD4 T cells while little to no chemotaxis (index >1) was detected from upper FRT CD4 T cells (**Fig. 4C & Fig. S5**). Next, we performed phenotyping of CD4 T cells by FRT compartment by measuring chemokine receptors and L-selectin expression previously identified from FRT T cells in humans (**Fig. 1&4D**). CD4 T cells from the lower FRT expressed greater CD62L (consistent with T_MM_) as compared with cervical and upper FRT CD4 T cells. FRT CD4 T cells from all tissue sites expressed a greater cell surface frequency of CCR5 and CCR6 as compared with lymph node memory CD4 T cells. Specifically, among FRT CD4 T cells, the lower FRT population was more likely to express CCR6 compared with upper FRT while lower FRT CD4 T cells were also more likely to express CCR5 compared with cervical-enriched CD4 T cells. As CX3CR1 expression in mice can distinguish memory T cells that perform tissue-itinerant immune surveillance functions (CX3CR1^int^) (*44*), we also assessed CX3CR1 expression from FRT CD4 T cells **(Fig. 4D&E).** Notably, CX3CR1^int^ CD4 T cells were most abundant in the lower FRT as compared with the cervical/upper FRT and as compared with LN memory CD4 T cells. Together, these data show that in mice FRT CD4 T cell populations comprise T_MM_ populations preferentially localized to the lower FRT and which display phenotypic properties consistent with migratory immune surveillance function.

**Figure 4.**
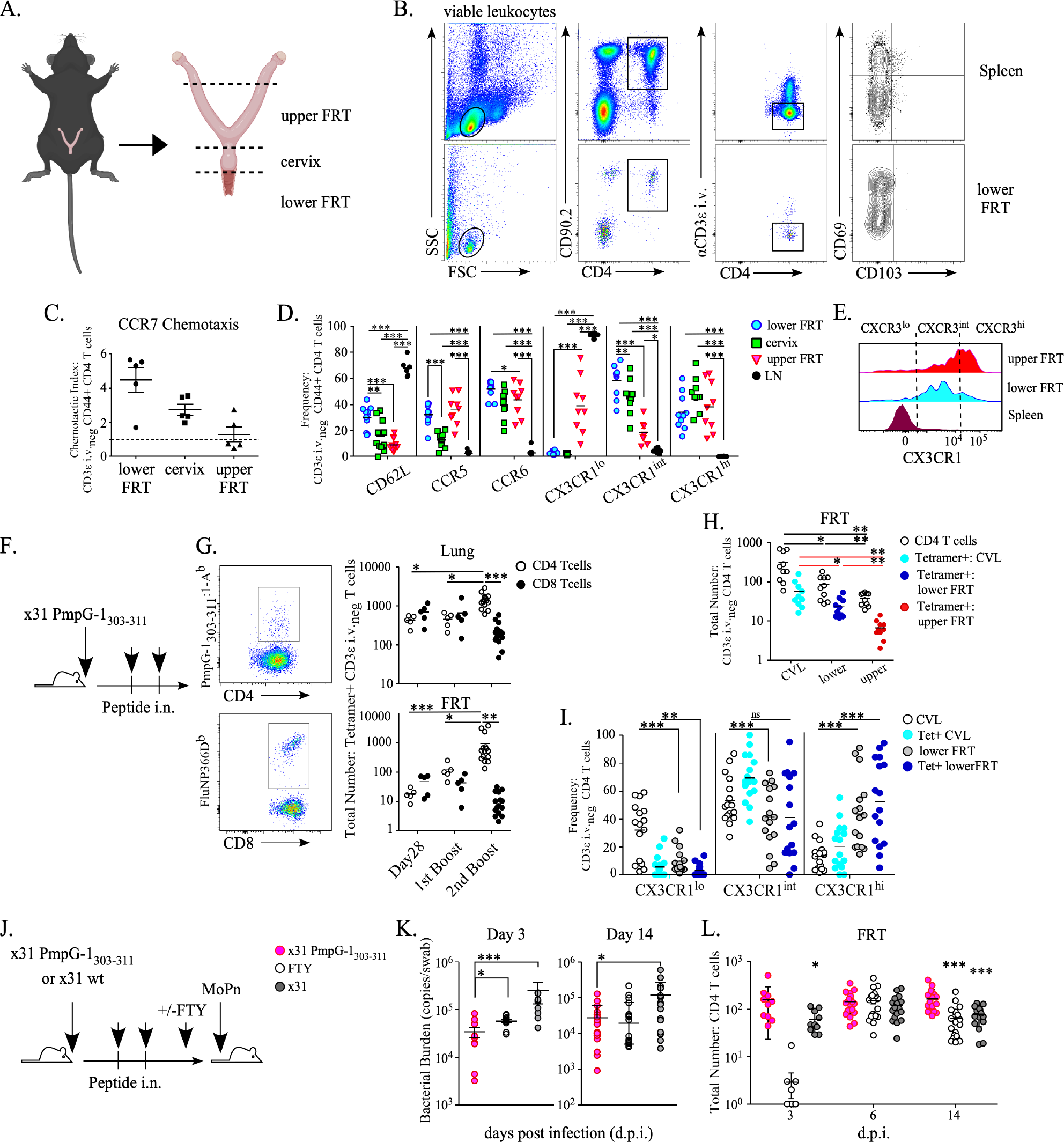
FRT CD4 T cell subset localization and response to *Chlamydia muridarum* challenge in mice. **(A).** Cartoon depiction of a C57Bl/6 mouse with FRT anatomic compartments indicated (created with BioRender.com). **(B).** Representative gating strategy for characterization of CD4 T cells resident in the FRT. **(C).** CD4 T cells enriched from the lower, cervical, or upper FRT region were measured for CCL21 chemotaxis via trans-well. The chemotactic index was calculated according to the absolute number of CD3 intravenous (IV) negative CD4 T cells migrating into CCL21 gradients and normalized to migration across trans-well into media alone. Graph represent 1 of 2 separate experiments (N=10). **(D).** The frequency of CD62L, and indicated chemokine receptors measured from IV negative CD4 T cells compared across FRT anatomic regions (N=10). **(E)**. Representative histograms demonstrating the cell gating strategy for CX3CR1 expression by mean fluorescent intensity (MFI) as compared among CD3 IV-negative CD4 T cells enriched from the upper FRT, lower FRT and IV+ CD4 T cells in circulation (spleen) to demonstrate distribution. **(F).** Experimental mouse model schematic for inducing *C. muridarum* probable outer membrane protein (PmpG-1)-specific CD4 T cells. **(G).** (Left panels) Representative staining of PmpG-1_303-311_-specific CD4 T cells (left, top) and influenza nucleoprotein (FluNP) specific CD8 T cells (left, bottom) by tetramer staining. (Right panels) the total number of CD3ε IV-negative PmpG-1_303-311_-specific CD4 (white circles) or FluNP-specific CD8 (black circles) T cells detected in the lung (top, right) or FRT (bottom, right) at 28 days post-infection with x31-PmpG-1 and following the first and second intranasal PmpG-1 peptide boost (N=10). **(H).** The total number of CD3ε IV-negative CD4 T cells (white circles) and PmpG-1_303-311_-specific CD4 T cells detected from CVL (light blue circles), lower FRT (dark blue circles), or upper FRT (red circles). **(I).** The CX3CR1 expression intensity of CD3ε IV-negative CD4 T cells (CVL represented by white circles and lower FRT represented by gray circles) and PmpG-1_303-311_-specific CD4 T cells in the CVL (light blue circles) or lower FRT (dark blue circles). **(J).** Experimental mouse model schematic illustrating the prime-boost strategy for inducing PmpG-1-specific CD4 T cell followed by vaginal challenge with *C. muridarum* (MoPn). **(J).** The mean bacterial burden detected from vaginal swabs at 3, 6, and 14 days post infection (dpi) from X31 PmpG-1 boosted mice (pink circles), x31 PmpG-1 boosted mice treated with FTY720 (day -1 to day 3 dpi) (white circles), and mice original infected with X31 with NP peptide boosting as a control (gray circle). Graph shown as mean value with standard error bars and connecting lines per group). **(K).** Bacterial burdens at 14 dpi shown as an aligned dot plot graph with mean and standard error bars. **(L).** The total number of CD4 T cells detected from CVL of mice from indicated groups at 3, 6, and 14 dpi. **(C-L).** Means evaluated by multiple comparisons *p≤0.05, **p<0.01, ***p<0.001, ****p<0.0001. Unless otherwise noted graphs are shown as scattered dot plots with mean line.

To investigate the contributions of FRT T_MM_ in immune defense, we used a previously reported murine model of *C. trachomatis* infection, *Chamydia muridarum* (*C. muridarum*) (*45*). For generating *C. muridarum*-specific CD4 T_MM_ without establishing *C. muridarum*-specific T_RM_ in the genital tract we developed an intranasal prime-boost strategy (**Fig. 4F**) using an x31 (H32N) influenza virus engineered to express the immunodominant I-A^b^ MHC class II epitope of the *C. muridarum* probable outer membrane protein epitope (PmpG-1_303-311_) (*45*). Mice were infected intranasally with x31-PmpG-1 followed by two intranasal PmpG-1_303-311_ peptide+CpG boosts to induce PmpG-1-specific CD4 T cells. Characterization of x31-PmpG-1 memory mice showed that PmpG-1 CD4 T cells were detected in both the lung and FRT, whereas FluNP-specific CD8 T cells generated during the initial x31-PmpG-1 infection were largely absent from the FRT (**Fig. 4G**). Within the FRT, PmpG-1 CD4 T cells primarily localized to the vaginal epithelial barrier (determined by CVL collection) with detectable but reduced populations in the lower FRT tissue and few cells detected in the upper FRT (**Fig. 4H**). As our previous T cell characterizations in mice identified CX3CR1^int^ as an indicator of T_MM_, we measured CX3CR1 from FRT PmpG-1 CD4 T cells (**Fig. 4I**). These data showed that tetramer+ CD4 T cells from CVL were predominantly CX3CR1^int^, while CX3CR1^hi^ CD4 T cells (consistent with T_EM_) were more likely to localize in the FRT tissues. Taken together these data show that T cell priming at the lung generates FRT T_MM_ surveillance predominantly to the lower FRT region.

Next, x31-PmpG-1 memory and control mice (intranasal infection with wild-type x31 followed by influenza NP _311-325_ peptide+CpG boosting), were challenged vaginally with 10^5^ i.f.u. (inclusion forming units) of *C. muridarum* (**Fig. 4J**). To determine the role of FRT T_MM_ trafficking in immune protection we also blocked cell egress from lymph nodes in a cohort of x31-PmpG-1 memory mice using daily FTY720 treatment one day prior to and 3 days following *C. muridarum* inoculation. PmpG-1 memory mice showed reduced *C. muridarum* burdens early in the infection course compared to control mice (3 days post infection [dpi]) (**Fig. 4K**). Notably, FTY720 treatment resulted in a significant increase in the bacterial burden indicating an important role for lymphocyte recruitment for optimal protection. In addition we observed more rapid bacterial clearance at 14 dpi when the peak T cell response is detected in unimmunized mice (*45*). Corresponding to the reduced bacterial burden, the number of FRT CD4 T cells in PmpG-1 memory mice was elevated at the FRT barrier (**Fig. 4L**). Together, these data show that antigen-specific T_MM_ function to provide an early and sustained response to *C. muridarum* at the FRT vaginal barrier which effectively reduces the bacterial burden.

### Systemic Th1-specific activation and FRT T_MM_ fluctuations occur over the menstrual cycle

During the luteal phase of the menstrual cycle, the decidualized endometrium utilizes pro-inflammatory immune signaling to recruit leukocytes (especially granulocytes) from the circulation in order to facilitate the tissue remodeling process driving menstruation(*46*). However, whether these periodic systemic and local immune events might also influence FRT migratory CD4 T cell surveillance are unclear. Therefore, we used the pig-tailed macaque model of human menstruation to profile CD4 T cell properties in the circulation and at the vaginal epithelium under menstrual cycle regulation(**Fig. 5A**)(*38*). From macaques, bi-weekly blood samples were collected from 6 cycling animals over 9 weeks in order to measure sex-hormones, cytokines/chemokines, and CD4 T cell properties over consecutive cycles.

**Figure 5.**
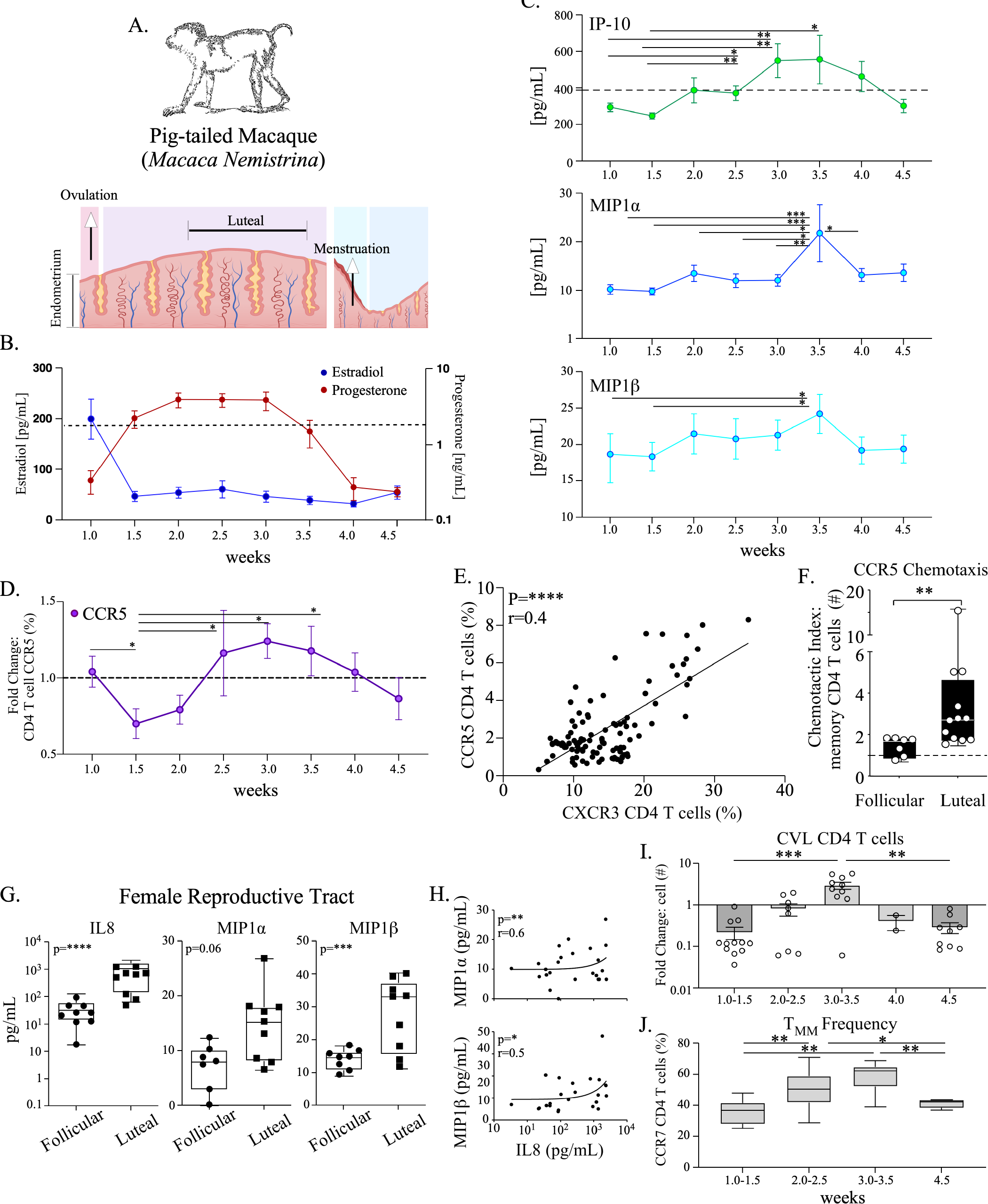
The luteal phase of the menstrual cycle is linked with increased CCR5 signaling and FRT T_MM_ frequency. **(A).** (Top) cartoon depiction of a pig-tailed macaque with (bottom) endometrial changes representing phases of the menstrual cycle. The time point of ovulation, the luteal phase (post implantation), and menstruation are indicated. Created with BioRender.com. **(B).** A mean line graph with standard error of the means (SEM) of P4 and E2 measured bi-weekly from six pigtail macaques are stratified by phase of the cycle at samples collection starting at the time of ovulation (designated week 1.0 and referenced to the cartoon depiction of cycle phase in A.). The average value of P4 is indicated by a dotted line for reference **(C).** A mean line graph with the SEM of indicated cytokine/chemokine concentrations measured from blood plasma are plotted over the cycle. **(D).** The fold change in cell surface CCR5 memory CD4 T cell expression frequency plotted over the cycle. The average value is indicated by dotted line. **(E)**. CCR5 frequency is correlated to CXCR3 from memory CD4 T cells and plotted as XY graphs with trend lines. Linear regression was evaluated using Pearson correlation. The Pearson correlation coefficients (r-value) and p-values (if ≤0.05) are shown **(F).** CCR5 chemotaxis from PBMC CD4 T cells collected at the follicular or luteal phase of the cycle and measured by trans-well assay. The chemotactic index was calculated by the absolute number of memory CD4 T cells migrating into Mip1α, Mip1β, and RANTES gradients and normalized to media controls. Shown as bar graph with SD bars overlaid with a scatter plot and mean line, dotted line represent chemotactic index of 1. Comparisons evaluated by Mann-Whitney test. **(G).** The concentration of IL-8, Mip1α, and Mip1β detected from CVL at the follicular or luteal phase of the menstrual cycle. **(H**). XY graph depicting the concentrations of IL-8 with Mip1α (top panel) or Mip1β (bottom panel). Correlations were evaluated by generalized multivariate regression modeling. Pearson correlation coefficients (r-value) and p-values (if ≤0.05) are shown. **(I)**. A bar graph overlaid with scatter dot plot and SEM depicting the fold change in CD4 T cell yields from CVL sampling over the cycle. Week 4.0 plots include samples collected prior to detection of menstruation onset. **(J)**. The frequency of CCR7^hi^ expression measured from FRT CD4 T cells over the cycle shown as box and whiskers graph with the SD and mean. Samples collected during the time frame of menstruation (week 4.0) are excluded **(I, J).** Means evaluated by multiple comparisons (**C-I**). *p≤0.05, **p<0.01, ***p<0.001, ****p<0.0001.

Progesterone [P4] and estradiol [E2] were measured from blood plasma to identify and stratify menstrual cycle phases for comparison (**Fig. 5B**). First, we investigated inflammatory cytokines/chemokines and observed elevations in Th1-associated IP-10, MIP-1α and MIP-1β (*47*) during the luteal phase (**Fig. 5C**). As CCR5-specific chemokine changes were identified, we next examined CCR5 expression on circulating memory CD4 T cell subsets over the cycle (**Fig. 5D-F**). By creating a kinetic profile of CCR5 expression frequency on CD4 T cells over time we identified cyclical changes over the course of a menstrual cycle, evidenced by significant elevations occurring during the luteal phase (**Fig. 5D**). To better define CCR5 increases we evaluated for correlations with memory CD4 T cell expression of CXCR3 as, in addition to CCR5, CXCR3 is a Th1 activation-associated chemokine receptor (*48*) (**Fig. 5E**). This analysis identified a positive linear relationship between the expression of CCR5 and CXCR3, suggesting an increased Th1 profile over the menstrual cycle consistent with previous reports (*21*). To investigate whether cycle phase was associated with trafficking of CD4 T cells to CCR5 chemokines, we compared CD4 T cell chemotaxis by trans-well assay at the follicular or luteal phase (**Fig. 5F**). This analysis showed that memory CD4 T cells were more likely to exhibit chemotaxis to CCR5-chemokines during the luteal phase. Taken together these results indicate that during the luteal phase, increased systemic pro-inflammatory signals are coupled with increased CCR5-specific trafficking by the CD4 T cells.

To evaluate whether corresponding changes in CCR5-signaling were also occurring in the FRT over the menstrual cycle, we measured vaginal lavage concentrations of CCR5 chemokines (MIP1α/β) in addition to IL-8, a chemokine upregulated in the FRT during the luteal phase of the menstrual cycle that functions to facilitate endometrial leukocyte recruitment into the endometrium (*49*). Weekly FRT sampling over the cycle showed higher MIP-1β levels during the luteal phase in parallel to IL-8 increases, whereas MIP1α, while also exhibiting an increased mean trend at the luteal phase, was not significantly different (**Fig. 5G and Supplementary Table 2**). To better define the relationship between IL-8 and CCR5 chemokines in the FRT, we performed generalized linear regression modeling on concentrations of IL-8 with MIP1α/β (**Fig. 5H**). These comparisons showed that IL-8 was positively correlated with both MIP1α and MIP1β, which suggested that FRT CCR5 chemokine production was similarly regulated by endometrial remodeling cues during the luteal phase of the menstrual cycle.

As we hypothesized that fluctuations in CCR5-signaling occur over the menstrual cycle might also influence migratory CD4 T cells surveillance in the FRT, we next asked whether T_MM_ populations were exhibiting fluctuations based upon cycle phase. To test this, we performed a phenotypic analysis from FRT CD4 T cell populations collected weekly by CVL and stratified according to basic cycle phases (**Fig. 5I**). By measuring the difference in CD4 T cell yields over time we identified increases occurring within the late luteal phase (identified as week 3.0-3.5) which then decreased prior to menstruation onset and remained reduced at the follicular phase (identified as week 4.5). Next, to determine the contribution of FRT T_MM_ to the differences in T cell yield we compared the frequency of FRT T_MM_ over the cycle (**Fig. 5J**). This analysis showed that the frequency of FRT T_MM_ increased from the timeframe of ovulation (identified as week 1.0-1.5) exhibiting peak mean levels at the late luteal phase which then decreased at the follicular phase to a level similar with those detected at ovulation. Taken together, these data show that consistent and predictable changes in both the circulating and FRT barrier CD4 T cell populations occur over the menstrual cycle, and that increased CCR5-specific signaling during the luteal phase is coupled with alterations in the composition of FRT CD4 T cell populations and most notably an increased frequency of T_MM_.

### CCR5 is required for steady-state memory CD4 T cell trafficking into the FRT following T_MM_ priming

Although CCR5-signaling has been previously reported as important for antigen-specific effector T cell trafficking into the FRT in response to local pathogen challenge in mouse models, the immune-mediated events of menstruation are not the result of infection. To first assess whether CCR5 signaling could also control CD4 T_MM_ migration into the FRT in the absence of a local infection, we used a mixed mixed bone marrow chimera approach in mice **(Fig. S6)**. WT (CD45.1^+^) and CCR5 KO (CD45.2^+^) mixed bone marrow chimeras were infected intranasally with x31-PmpG-1 and boosted with PmpG-1_303-311_ peptide+CpG to induce FRT memory CD4 T cell trafficking (**Fig. S6A**). We observed that WT CD4 T cells, including tetramer+ cells, largely outnumbered the CCR5 KO CD4 T cells in both the CVL and FRT as compared to the lung and spleen. The difference in WT compared with CCR5 KO was most pronounced in CVL with little to no detection of CCR5 KO cells. Taken together, these data demonstrate that CCR5 regulates memory CD4 T cell access to the FRT following priming at a distal tissue site.

### Maraviroc inhibits FRT T_MM_ specific migration during the luteal phase

Our previous results had so far evidenced that the composition of the FRT CD4 T cell population was changing over the course of the menstrual cycle and chemokines associated with CCR5-mediated trafficking were elevated during the luteal phase. Thus, to determine whether the menstrual cycle regulates CCR5-mediated trafficking and migration of T_MM_ into the vaginal barrier, we administered an oral CCR5 antagonist drug, Maraviroc (MVC), daily for a course of five days in pig-tailed macaques during either the follicular or luteal phase of the menstrual cycle (**Fig. S7 6A-B**). To asses drug penetrance in the genital tract based upon phase of the menstrual cycle, MVC drug levels were measured from CVL supernatants before (day 0) and following 5 days of MVC treatment (**Fig. 6C**). By day 5 of treatment, MVC concentrations were detected at the vaginal barrier during both the follicular and luteal phases of the cycle. The efficacy of MVC binding to CCR5 was assessed by measuring receptor occupancy, and both T_CM_ and T_EM_ CD4 T cells from PBMC confirmed MVC action through a reduction in CCR5 internalization at day 5 of MVC treatment (**Fig. 6D-E**).

**Figure 6.**
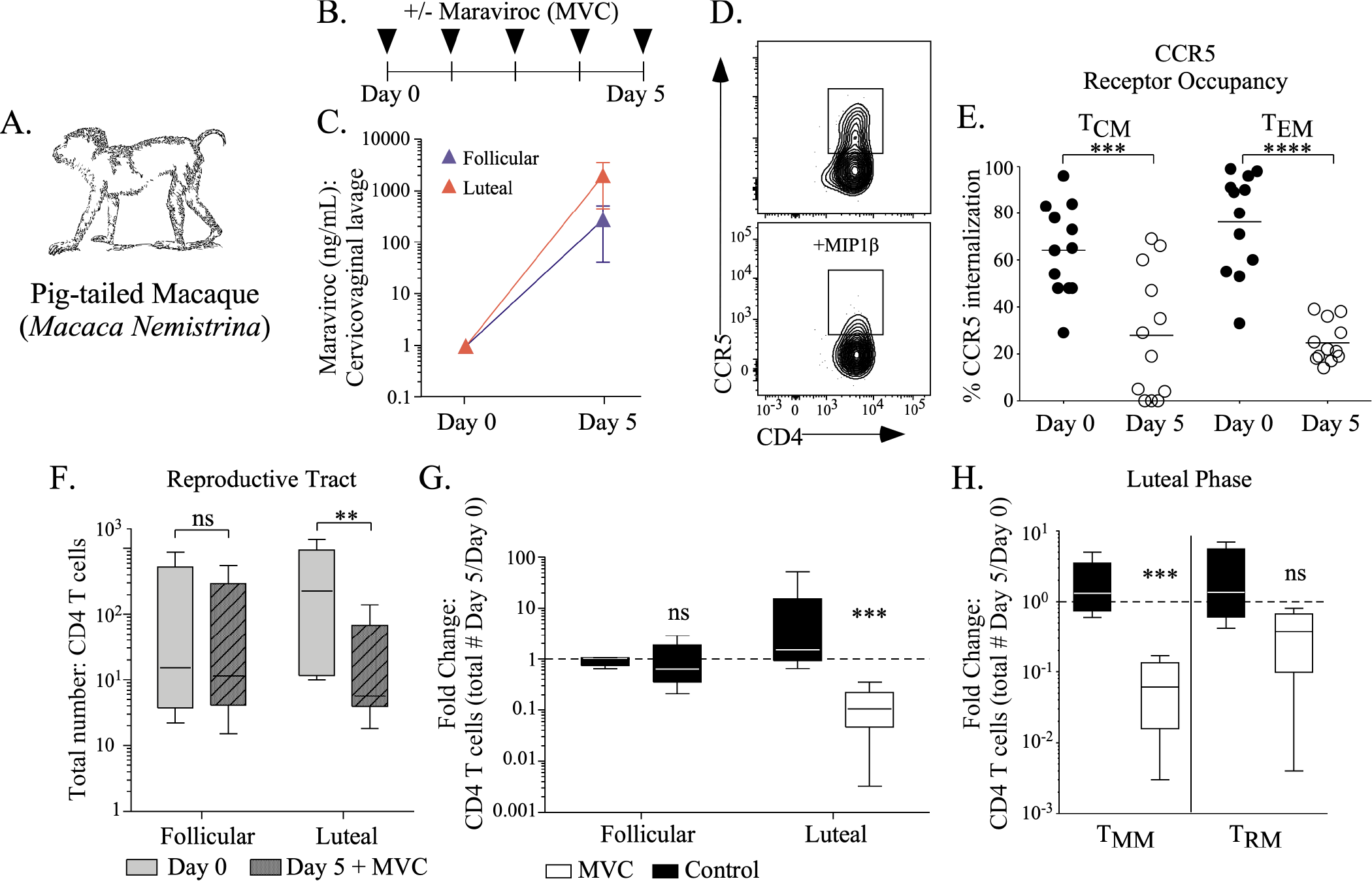
Oral Maraviroc inhibits migration of T_MM_ to the FRT during the luteal phase. **(A).** Schematic diagram of oral Maraviroc (MVC) treatment in six pigtail macaques. MVC was administered daily for 5 days. **(B).** MVC drug levels detected in CVL collected at either the follicular or luteal phase of the menstrual cycle at indicated time points prior to (Day 0) or following five days of oral MVC treatment (Day 5). **(C).** Representative flow plots depicting the receptor occupancy assay for CCR5 internalization. CD4 T cells from PBMC were measured for cell surface CCR5 expression (top panel) with or without incubation with MIP1β (bottom panel). **(D).** CCR5 internalization from either CCR7^hi^ or CCR7^lo^ memory CD4 T cells from PBMC measured from animals prior to (black circles) or following five days of MVC treatment (white circles). Means were evaluated by multiple comparisons. Graph shown as scatter dot plot. *p≤0.05, **p<0.01, ***p<0.001, ****p<0.0001**. (E).** The absolute number of CVL CD4 T cells were measured before (Day 0) and following a 5-day course of MVC at either the follicular or luteal phase of the menstrual cycle. **(F).** The fold change of the absolute number of CVL CD4 T cell comparing drug treatment (white box) to internal placebo control (black box). **(G).** Fold change of the absolute number of CVL CD4 T cells compared by CCR7 expression. **(E-G)** Graphs depicted as box and whiskers plot indicating the SD and mean. Models used to evaluate means were determined by linear regression and fit using generalized estimating equations (GEE). p-values ≤0.05 are shown.

To measure the impact of MVC on the memory CD4 T cell population in the lower FRT, the number of total FRT memory CD4 T cells was determined before and after drug treatment. While no change in the number of memory CD4 T cells was detected from MVC-treated animals during the follicular phase, a significant reduction of memory CD4 T cells was detected in MVC-treated animals during the luteal phase (**Fig. 6F**, **Fig. S7**). Similarly, by analyzing the fold change (from the estimated absolute count of FRT CD4 T cells detected at day 5 normalized to day 0 for each animal) comparing MVC treatment to placebo using a multiple linear regression estimation model, a reduction in memory CD4 T cells was detected with MVC treatment during the luteal phase of the menstrual cycle (**Fig. 6G**). Notably, we observed a specific reduction within the CD4 T_MM_ population, whereas no difference was detected with CD4 T_RM_ following MVC treatment during the luteal phase (**Fig.6H**). Taken together, these data illustrate that during the luteal phase of the menstrual cycle, increased CCR5 signaling directs T_MM_- specific trafficking into the lower FRT barrier.

## Discussion

Memory T cells in the FRT are essential for optimal protection against a variety of genital tract infections (*50*). Although the vast majority of women and adolescent girls of reproductive age experience spontaneous decidualization and menstruation periodically over a lunar revolution (*51*), the impact of menstruation on the composition of the FRT memory CD4 T cell population and T cell-mediated protection against STIs remains poorly understood (*52*). Here, we report that the FRT memory CD4 T cell pool is highly dynamic and comprised of both T_MM_ and T_RM_ subsets in humans, primates, and mice. Notably, localized inflammation and tissue remodeling during the luteal phase of the menstrual cycle results in a transient but profound shift in the ratio of T_MM_: T_RM_ subsets in the FRT driven by the CCR5-mediated recruitment of T_MM_ from the circulation. Thus, unlike memory CD4 T cell populations in other barrier tissues, the FRT memory CD4 T cell pool undergoes substantial and repetitive fluctuations in the absence of a local infection, due to the process of menstruation.

Previous studies of memory CD8 T cell subsets in peripheral tissues have shown these populations to be primarily comprised of T_RM_ that permanently reside in the tissue and T_EM_ that are transiting through the tissue while conducting immunosurveillance (*40, 53*). Both of these memory CD8 T cell subsets are characterized by a lack of the lymph node homing receptor CCR7, in addition to expression of tissue-retention molecules within the T_RM_ subset (*53*). While tissue memory CD4 T cell populations are also comprised of these subsets, studies in the skin of mice and humans identified an additional CD4 T cell subset, T_MM_, that expresses CCR7 (*25, 32*). In the FRT, we found that the memory CD4 T cell population was enriched for CCR7^+^ T_MM_. Consistent with previous reports from skin, FRT T_MM_ expressed molecules associated with migration to non-lymphoid tissues and sites of inflammation.

Notably, we observed that FRT CD4 T_MM_ in mice exhibited intermediate expression of CX3CR1, which is associated with a distinct population of memory CD8 T cells specialized for surveillance of peripheral tissues (*42*). Although it is unclear whether a similar function can be attributed to memory CD4 T cells expressing intermediate levels of CX3CR1, it is consistent with observations of T_MM_ surveilling both peripheral and lymphoid tissues. Finally, unlike FRT T_RM,_ FRT T_MM_ were primarily localized to the lower FRT. One potential explanation for this compartmentalization is the differential composition of the epithelial barrier in the lower versus upper FRT, with the lower FRT barrier being comprised of a stratified epithelial layer similar to the skin that allows for trafficking of memory CD4 T cells between the circulation and tissue under steady-state conditions (*54*). Together, these findings demonstrate that the memory CD4 T cell pool in the lower FRT is a heterogenous population comprised of both tissue-anchored T_RM_ and transiting T_MM_ capable of migrating from the tissue back to lymphoid sites.

The ontogeny of T_MM_, and their relatedness to T_RM_ or T_CM_, is still an open area of investigation. The ability of T_MM_ to traffic to lymphoid tissues in a CCR7-dependent manner closely aligns with T_CM_, and our observation of increased IL-2 production by T_MM_ in comparison to T_RM_ suggests effector functions more similar to T_CM_ (*55*). In contrast, the broad expression of chemokine receptors and adhesion molecules on T_MM_, and their localization in the FRT, are more closely aligned with T_RM_. We found that RNA sequencing of FRT T_RM_, FRT T_MM_, and lymph node T_CM_ clearly distinguish T_MM_ from T_CM_, and that T_MM_ are transcriptionally similar to T_RM_. However, it is important to note that genes associated with the core transcriptional program upregulated during T_RM_ development (*41*) are not enriched in T_MM_, suggesting that T_MM_ are not merely a subset of T_RM_ that express CCR7. We and others have previously shown that tissue entry can drive substantial transcriptional re-programming of memory T cells, irrespective of the core T_RM_ program, as T cells adapt to the local microenvironment (*56, 57*). Thus, the similar transcriptional profiles of FRT T_MM_ and T_RM_ could be driven, at least in part, by their co-localization in the tissue.

CCR5 chemokines are constitutively produced from FRT epithelial cells, local innate immune cells and potentially by underlying stromal cells (*8, 14, 22, 58, 59*). Furthermore, artificially modifying FRT barrier integrity through high dose progestin-based contraception such as injectable depot medroxyprogesterone acetate (DMPA) leads to an increase in FRT CCR5-expressing CD4 T cells (*60, 61*), suggesting that progesterone signaling pathways can modulate CD4 T cell trafficking to the FRT through changes in the vaginal epithelium or local chemokine production. Our data over the course of the menstrual cycle support this premise as the primary mechanism for T_MM_ migration to the lower FRT, as CCR5-deficient antigen-specific CD4 T_MM_ generated by a respiratory infection were severely impaired in their ability to migrate to the FRT in mice. Furthermore, analysis of chemokine production in the FRT of NHP showed a significant increase in CCR5-binding chemokines during the luteal phase of the menstrual cycle, which correlated with increased systemic progesterone levels. Finally, blocking CCR5 signaling in NHP at the luteal, but not follicular, phase of the menstrual cycle resulted in a significant decrease in the number of CD4 T_MM_ in the lower FRT. These data demonstrate an essential role for CCR5 signaling in the migration of T_MM_ to the FRT, especially at the luteal phase when local production of CCR5-binding chemokines is increased.

One interesting area of future investigation will be to determine whether T_MM_ can convert into T_RM_ within the tissue if given the proper signals. Previous studies have shown that entry of effector T cells into some peripheral tissues, combined with signaling through the TGF-β receptor, was sufficient to promote their differentiation to T_RM_ (*62*). This raises the possibility that the periodic influx of T_MM_ into the FRT during the luteal phase of the menstrual cycle could lead to a shifting FRT CD4 T_RM_ population over time as T_MM_ convert to T_RM_. Our also data demonstrate that CD4 T_MM_ surveillance of the FRT is important for cell-mediated protection against a genital tract infection, as preventing antigen-specific CD4 T_MM_ entry into the FRT by treatment with FTY720 resulted in increased bacterial loads following *C. muridarum* infection. It is tempting to speculate that increased trafficking of T_MM_ to the FRT during the luteal phase of the cycle may be a mechanism to increase the local antigen-specific CD4 T cell repertoire at a time when tissue remodeling can influence the integrity of the epithelial barrier. Future experiments are needed to fully define the roles of pathogen-specific T_MM_ in protection against sexually transmitted infections, and to determine the potential for T_MM_ to differentiate into T_RM_ in the FRT, especially in relation to the phases of the menstrual cycle.

To conclude, this study finds that cellular immune surveillance in the FRT is regulated through the local production of CCR5-binding chemokines, which direct the recruitment of a unique population of memory CD4 T cells, T_MM_, that are poised to migrate between tissues and lymphoid organs. The menstrual cycle regulates the magnitude of T_MM_ migration to the FRT through periodic oscillations in chemokine production that are increased at the luteal phase of the cycle, correlating with progesterone levels. These discoveries underscore the need for further investigations into understanding menstrual cycle regulation of cellular immune memory in the FRT and how this process can impact both immunity against STIs and infection-risk in women.

## Materials and Methods

### Human specimens

Human paired blood and CVL specimens were obtained from women participating in a study assessing the effects of hormonal contraception on HIV biomarkers prior to starting hormonal contraception at Grady Clinic (Atlanta, Georgia). Women interested in initiating a new contraceptive were recruited into the study through community referral or self-referral and were screened for study eligibility, including HIV-negative status (OraQuick swab) and no recent STIs (self-report and later tested by reverse transcription PCR described below). Upon enrollment, participants were scheduled for their first study visit (baseline) approximately 3 weeks after their LMP next menstrual period, corresponding with their luteal phase. Additional eligibility for participation required that women were aged 18–45 years, exhibiting normal menses (22-to 35-day cycles for at least 3 cycles), not using any hormonal contraception method or intrauterine device for at least 6 months prior, not pregnant for at least 3 months prior, not breastfeeding, and not experiencing symptomatic vaginal infection or genital ulcer disease at screening. Participants did not have a history of loop electrosurgical excision procedure, conization, or cryosurgery in the previous year. Participants were further screened for medical eligibility based on the Centers for Disease Control and Prevention (CDC) medical eligibility criteria for contraceptive use and clinical judgment (*63*). Protocols were approved by the Emory University and CDC Institutional Review Boards and the Grady Research Oversight Committee. All participants provided informed consent.

### Sexually Transmitted Infection Testing

DNA was extracted from DrySwab^®^ (Lakewood Biochemical Company) using the Qiagen DNA Mini kit and used to amplify targets from *Neisseria gonorrhoeae*, *Chlamydia trachomatis*, Herpes simplex virus types 1 and 2, *mycoplasma genitalium*, and *Trichomoniasis vaginalis*, using real-time duplex PCR and Qiagen Rotor-Gene Q and 6000 real-time PCR instruments. Qiagen Rotor-Gene Q Series software was used to analyze data.

### NHP Ethics statement

All of the animal procedures performed in the present study were approved by the Institutional Centers for Disease Control and Prevention (CDC) Animal Care and Use Committee. Macaques were housed at the CDC under the full care of CDC veterinarians in accordance with the standards incorporated in the *Guide for the Care and Use of Laboratory Animals* (National Research Council of the National Academies, 2010). SHIV-infected macaques or uninfected macaques in moribund decline were humanely euthanized in accordance with the American Veterinary Medical Association Guidelines on Euthanasia, June 2007. All procedures were performed under anesthesia using ketamine, and all efforts were made to minimize suffering, improve housing conditions, and provide enrichment opportunities.

### NHP specimens

A total of 16 pig-tailed macaques were used. For animals administered MVC, drug preparation and administration were carried out as previously described (*64*). During time points of MVC treatment drug was administered orally by gavage once daily for a course of five days. 3 adult female rhesus macaques were used to study a SHIV_162P3_ recall response from CD4 T cells. Animal were challenged rectally with 10TCID_50_ SHIV_162P3_ weekly until infection was detected by RT-PCR from blood plasma. SHIV RNA was measured from blood plasma weekly using a modified one-step reverse transcriptase PCR (RT-PCR) method as previously described (15 minutes at 48^0^C sensitivity 50 RNA copies/ml) (*65*). The estimated time of infection was defined as 7 days prior to the first positive RNA RT-PCR in plasma to account for the eclipse period between virus inoculation and detection of SHIV RNA in plasma.

Animals were euthanized between 20-23 weeks post infection.

### Intravascular labeling in NHP

Intravascular staining in macaques was performed as previously described (*39*). Briefly αCD45 (BD Biosciences) conjugated to R-phycoerythrin (PE) was diluted into sterile saline and IV injected into the saphenous vein and allowed to circulate for five minutes prior to tissue collection.

### Microscopy

Tissues were flash-frozen in OCT by floating on liquid nitrogen. Blocks were sectioned at 7 microns on a cryostat, and slides were fixed in 75:25 acetone/ethanol. The slides were blocked with FACS wash containing 1 μg/ml anti-mouse CD16/32 (clone 2.4G2), 10% mouse serum, 10% rat serum, and 10% donkey serum. Antibodies were from Abcam: rabbit anti-pan cadherin Alexa Fluor® 488 (clone EPR1792Y), rabbit anti-CCR7 Alexa Fluor® 488 (clone Y59), with DAPI staining solution; BioLegend: anti-mouse TCR α/β -Alexa Fluor® 647 (clone R73). Coverslips with Prolong Gold were applied, and the slides were cured overnight before imaging. Imaging was performed on a Zeiss Axio Observer Z1 with an Axiocam 506 monochromatic camera. Image processing was performed with Zen 2 software.

### RNA sequencing

For each population, 1,000 cells were sorted into RLT lysis buffer (Qiagen) containing 1% BME and total RNA purified using the Quick-RNA Microprep kit (Zymo Research). All resulting RNA was used as an input for complementary DNA synthesis using the SMART-Seq v4 kit (Takara Bio) and 10 cycles of PCR amplification. Next, 1 ng cDNA was converted to a sequencing library using the NexteraXT DNA Library Prep Kit and NexteraXT indexing primers (Illumina) with 10 additional cycles of PCR. Final libraries were pooled at equimolar ratios and sequenced on a HiSeq2500 using 50-bp paired-end sequencing or a NextSeq500 using 75-bp paired-end sequencing. Raw fastq files were mapped to the MacamM v7 build of the Rhesus macaque genome (*66*) using STAR (*67*) with the v7.8.2. transcriptome annotation library reference transcriptome (*66*). PCR duplicate reads were flagged with PICARD mark duplicates and excluded from downstream analyses. The overlap of reads with exons was computed and summarized using the GenomicRanges (*68*) package in R/Bioconductor and data normalized to fragments per kilobase per million (FPKM). Genes that were expressed at a minimum of three reads per million (RPM) in all samples for each cell typed were considered to be expressed. DEGs were determined using the glm function in edgeR (*69*) using the animal from which each cell type originated as a covariate. Genes with an FDR < 0.05 and absolute log_2_(FC) > 1 were considered to be significant. For GSEA (*70*), all detected genes were ranked by multiplying the sign of the fold change by the −log_10_ of the *P* value between two cell types. The resulting list was used in a GSEA pre-ranked analysis. All code for data processing and display is available upon request.

### Tissue specimen processing

Human and Non-human primate samples were processed similarly: Blood was collected in 8 mL sodium citrate-containing CPT tubes (BD Biosciences) and separated into plasma and peripheral blood mononuclear cells (PBMC) by centrifugation. CVL specimens were collected and leukocytes enriched as previously described (*24*). CVL samples with blood detection (naïve T cells and B cells) were excluded from analysis. NHP FRT tissues were further dissected and processed into single cell suspensions using collagenase type II (62.5 U/ml), and DNase I (0.083 U/ml). Cell suspensions were separated by Percoll (GE healthcare life sciences) discontinuous density centrifugation. Enriched cell populations were washed and resuspended in media for phenotyping or functional assays.

### Sex-hormone measurement and estimating menstrual cycle phase

Progesterone [P4] and Estradiol [E2] levels in plasma were quantified by immunoassay as one batch. Assay services were provided by the Biomarkers Core Laboratory at the Yerkes National Primate Research Center. Menstrual cycle phase was estimated by P4 and E2 kinetics relative to a 32-day menstrual cycle (average length of pigtail macaque menstrual cycle) and by observed menstruation.

### Soluble Cytokine/Chemokine Measurement

CVL supernatant and blood plasma was measured and analyzed for cytokines/chemokines through the Emory Multiplexed Immunoassay Core using the Meso Scale discovery (MSD) platform with NHP multiplex kits as one batch.

### Maraviroc drug measurement

The concentrations of Maraviroc (MVC) in mucosal secretions were measured by HPLC-MS/MS (Shimadzu Scientific, Sciex,). Briefly, MVC was extracted from Weck-Cel surgical spears (Beaver-Visitec International, Inc.) used to collect vaginal secretions (∼25 μl), using 500 μl of methanol containing internal standard (deuterium-labeled MVC [MVC-d6]; Toronto Research Chemicals). After a brief centrifugation the extracted liquid was evaporated to near dryness and then re-suspended in 150 μl of mobile phase A (0.2% formic acid in water). A 10-μl portion of the final solution was injected into a UK-C18 column (1 x 100 mm; Imtakt) connected to the HPLC-MS/MS system. An aqueous-acetonitrile mobile-phase gradient was used to elute the MVC from the column and into the MS/MS system. MVC mass transitions (*m/z*) of 514.1/117.2 and 514.1/389.5 were monitored in positive MRM mode. MVC concentrations were calculated using Analyst software (Sciex) from a standard curve with a dynamic range of 1 to 2,000 ng/ml. The lower limit of quantification of this assay was 5 ng/ml.

### Analysis of MVC binding to CCR5 by a MIP-1β internalization assay

MVC binding to CCR5 was determined *ex vivo* using a MIP-1β internalization assay as previously described (*64*). In brief, 1×10^6^ PMBCs were incubated with 20 ng of MIP-1β (recombinant human CCL4L1/MIP-1β isoform LAG-1; R&D Systems) for 30 min at 37°C. Cell surface CCR5 receptors on CD4 T cells are internalized upon binding to MIP-1β (a natural ligand of CCR5) (*71*) and MVC binding to CCR5 prevents internalization by MIP-1β, allowing extracellular CCR5 measurement by flow cytometry.

### Mice

C57BL/6J (wild-type [WT]), B6.SJL-*Ptprc^a^ Pepc^b^*/BoyJ (CD45.1), and B6.129P2-*Ccr5^tm1Kuz^*/J (CCR5 KO) mice from The Jackson Laboratory were housed under specific Animal Biosafety Level 2 conditions at Emory University. All experiments were completed in accordance with the Institutional Animal Care and Use Committee guidelines of Emory University.

### Intravascular staining in mice

Intravascular staining in mice was performed prior to euthanasia and tissue harvest as previously described (*43*). In brief, to identify T cells resident in various tissues, 1.5μg fluorophore-conjugated anti-CD3ε Ab in 200 μl 1× PBS was IV-injected into the tail vein of mice; 5 minutes post-injection, mice were euthanized with Avertin (2,2,2-tribromoethanol; Sigma-Aldrich) and exsanguinated prior to CVL and tissues collection.

### Mixed bone marrow chimeras

Mixed bone marrow chimeras were generated as previously described (*72*). Briefly, bone marrow cells were harvested from the femurs of naïve WT (CD45.1) and CCR5KO (CD45.2) donor mice, and a 1:1 mixture was then transferred intravenously into lethally irradiated (950 rads) CD45.1/CD45.2 recipient mice. Recipient mice were rested for 8 weeks to allow for reconstitution, and a blood sample was taken to confirm chimerism prior to use in experiments.

### Recombinant influenza virus and prime boost regimen

Using commercial gene synthesis, the nucleotide sequence of the PmpG-1_300-315_ peptide (SPIYVDPAAAGGQPPA: 5’ TCT CCA ATT TAT GTG GAC CCA GCT GCT GCA GGA GGA CAA CCT CCA GCT 3’) was inserted into the x31 influenza A virus neuraminidase (NA) gene at the site corresponding to position 44-45 of the NA stalk domain. This peptide encompasses the core I-A^b^ CD4 T cell epitope PmpG-1_303-311_. The modified NA gene was then cloned into the pDP reverse genetics vector (*73*) and combined with plasmids encoding the remaining seven gene segments of x31 for reverse genetics-based recovery of x31-PmpG-1 virus, as previously described (*74*). A working stock of x31-PmpG-1 virus was generated by inoculating 11 day old embryonated chicken’s eggs with transfected 293T cells plus associated culture medium from the reverse genetics procedures. Eggs were incubated for 40 h at 37°C prior to chilling and harvesting of allantoic fluid. This fluid was aliquoted and stored at -80°C. Insertion of the PmpG-1 was confirmed by sequencing of viral cDNA. Mice were infected intranasally with 3×10^4^ PFU of x31-PmpG-1 virus and boosted intranasally with 1μg PmpG-1_303-311_ peptide + 2μg CpG (ODN 1826, Invivogen) on days 14 and 28 (*75*). Mice were rested with 28-35 days following the final boost prior to analysis or challenge with *C. muridarum*.

### Chlamydia infection

*Chlamydia muridarum* (*C. muridarum*) Mouse Pneumonitis NiggII strain (ATCC^®^) was cultured in HeLa cells and purified by density centrifugation as previously described (*45*) aliquots were stored in sodium phosphate glutamate buffer (SPG) at -80C. The inclusion forming units (IFUs) from purified elementary bodies (EB) was determined by infection of HeLa 229 cells and enumeration of inclusions by microscopy. For vaginal infection 10^5^ IFU of *C. muridarum* in 10 µL SPG buffer were deposited into the vaginal vault. To measure bacterial burden, DNA was purified from vaginal swabs (Qiagen) and quantitated by ddPCR.

### ddPCR

The *C. muridarum* bacterial burden was measured using Droplet Digital™ PCR (ddPCR™) technology (Bio-Rad) according to manufacturer recommendations. In brief, a mixture containing 11uL of 2x QX200™ ddPCR™ EVAgreen® supermix, 2.2uL of mixed 16SR (chlamydia muridarum) forward (AGTCTGCAACTCGACTAC) and reverse (GGCTACCTTGTTACGACT) primers (4 uM), 4.4uL of ultrapure water, and 4.4 uL of cDNA sample was used to amplify a fragment of the gene of interest. 20 uL of this mixture was added to 70uL of droplet generation oil, and after the droplet generation step, 40 uL of the suspension was used to perform ddPCR in a 96-well PCR plate. The fluorescent signal was read by a QX200™ Droplet Reader (BioRad) and analyzed with QuantaSoft software. The gating for positive droplets was set according to the positive and negative controls read with each plate.

### Chemotaxis assay

Chemotactic activity was measured using a standard migration assay (*76*). In brief, either RANTES + MIP1β (300ng/mL) or CCL21 (300ng/mL) chemokines (R&D Systems, Minneapolis, Minnesota) were added to the lower chamber of 24-well transwell plates (5.0 μm pore size, Corning, Tewksbury, Massachusetts) and incubated 20 minutes at 37°C. Cells (up to 1 x 10^6^) were added to the upper chamber and incubated 3 hours at 37°C. Lower chamber contents were analyzed by flow cytometry as indicated using 123 count eBeads (affymetrix eBioscience) to calculate the absolute number of cells.

Chemotactic index was calculated as the number of cells migrating in response to indicated chemokines divided by the number of cells migrating in response to media alone.

### Statistical Analysis

#### Generalized estimating equations (GEE)

In the event of comparisons with repeated measures, models used to compare cellular immune markers and levels of soluble inflammatory cytokines concentrations by cycle phase were fit using generalized estimating equations (GEE) with an exchangeable working correlation structure.

#### Regression modeling

Correlations of IL-8 with CCR5 chemokines in Macaques were evaluated by generalized multivariate regression modeling to adjust for the subject effect followed by calculation of Pearson correlation coefficients on the residuals.

#### Regression model for cellular characteristics following oral MVC administration in macaques

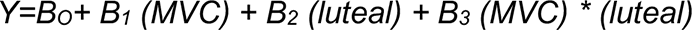

Absolute counts of CD4^+^, CD4^+^ CCR7^+^, and CD4^+^ CCR7^-^ T cells recorded on the day when treatment was given (Day 0) and 5 days after treatment (Day 5). The outcome we used in the analysis was the log-transformed ratios of cell counts or percentages on Day 0 to Day 5. Samples collected during menstruation were excluded (CD). The collected data are summarized in the table below.

| <u>Phases</u><br><u>Treatment</u> | <u>Follicular</u><br><u>MVC</u> | <u>Follicular</u><br><u>placebo</u> | <u>Luteal</u><br><u>MVC</u> | <u>Luteal</u><br><u>placebo</u> |
| --- | --- | --- | --- | --- |
| 1 | 1 | 1 | 1 | 1 |
| 2 | 1 | CD | CD | 1 |
| 3 | 2 | 1 | - | 1 |
| 4 | - | 1 | 2 | 1 |
| 5 | 1 | 1 | 1 | 1 |
| 6 | - | CD | 2 | 1 |

As a longitudinal unbalanced design, we used a multiple linear regression with generalized estimating equations (GEE) to adjust correlations between outcomes for the same subject. Binary variables MVC (1 if MVC was given and 0 for placebo) and luteal (1 if treatment was administered in luteal phase and 0 for follicular phase), and their interactions were included in the regression model.

Unless otherwise noted a paired two-tailed Student t-test was performed with Prism 5 software (Graphpad) to determine significance: *p<0.05, **p<0.01, ***p<0.001, ****p<0.0001.

### Intracellular Cytokine Staining

Cells in cRPMI-10 with 20 U/ml IL-2 (Roche) were activated with Cell Stimulation Cocktail (Affymetrix eBioscience) or overlapping SHIV_162P3_ envelope peptides for 5 hours (AIDS Research and Reference Reagent Program, Division of AIDS, National Institute of Allergy and Infectious Diseases, National Institutes of Health). Following surface staining, cells were permeabilized using a Cytofix/Cytoperm kit as recommended by the manufacturer (BD Biosciences) prior to intracellular antibody staining Background values from matched untreated populations were subtracted from cytokine measurements.

### Cell staining and flow cytometry

Cells enriched from various tissues were stained for viability using the Zombie Fixable Viability Kit (Biolegend^®^) then incubated with anti-CD16/32 Fc block (BioXcell). Tetramers were provided by the National Institutes of Health (NIH) Tetramer Core Facility at Emory University, class I H-2D^b^ Influenza A NP 366-374 (ASNENMETM) conjugated to Allophycocyanin (APC) and class II I-A^b^ C. murid pmpG-1 303-311 (YVDPAAAGG) conjugated to R-phycoerythrin (PE). Tetramer staining occurred prior to cell surface staining. Stained samples were run on a LSRII flow cytometer or NHP cells mechanically sorted using a Sony SH800 and data were acquired using FACS DIVA software (BD Biosciences) and analyzed using FlowJo software (TreeStar, Inc.). The following fluorochrome conjugated antibodies were used:

### Human antibodies

**Human antibodies:**
| antibody | conjugate | clone | vendor |
| --- | --- | --- | --- |
| CD3 | Brilliant Violet 421™ | UCHT1 | Biolegend® |
| CD4 | Alexa Fluor® 700 | RPA-T4 | BD Biosciences |
| CD4 | Brilliant Violet 785™ | RPA-T4 | Biolegend® |
| CD8a | Brilliant Violet 510™ | RPA-T8 | Biolegend® |
| CD45 | APC/FIRE™750 | 2D1 | Biolegend® |
| CD45RA | Brilliant Violet 605™ | HI100 | Biolegend® |
| CCR7 | PE-CF594 | 150503 | BD Biosciences |
| CXCR3 | PerCp/Cyanine5.5 | G025H7 | Biolegend® |
| CX3CR1 | PerCp-eFlour®780 | 2A9-1 | affymetrix eBioscience |
| CCR4 | PerCP/Cyanine5.5 | IG1 | Biolegend® |
| CCR5 | PE | 3A9 | BD Biosciences |
| CCR6 | Brilliant Violet 605™ | G034E3 | Biolegend® |
| CCR9 | APC | L053E8 | Biolegend® |
| *α4β7 |  |  | NIH AIDS reagent program |
| IFNγ | APC | 25723.11 | BD Biosciences |
| IL-2 | Brilliant Violet 605™ | MQ1-17H12 | BD Biosciences |
| TNFα | Brilliant Violet 711™ | Mab11 | Biolegend® |
| IL-17A | PE | N49-653 | BD Biosciences |
| CD62P | PerCp-eFlour®780 | Psel.KO2.3 | affymetrix eBioscience |
| CD62L | Brilliant Violet 650™ | DREG-56 | Biolegend® |
| CD66b | APC | G10F5 | BD Biosciences |
| CD69 | Brilliant Violet 650™ | FN50 | Biolegend® |
| CD103 | fluorescein isothiocyanate | Ber-ACT8 | BD Biosciences |

\*conjugated to APC (Molecular probes).

### NHP antibodies

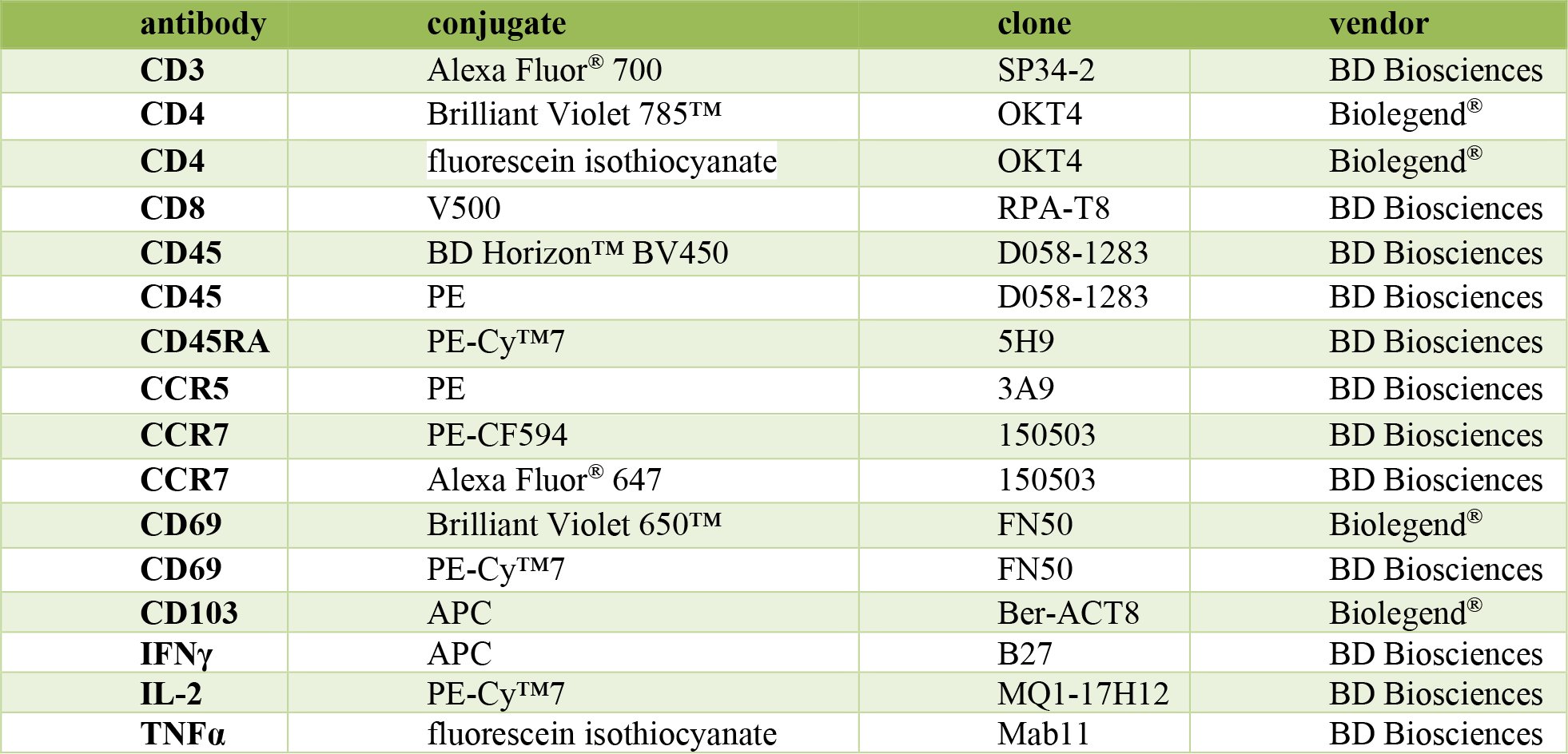

NHP antibodies:
| antibody | conjugate | clone | vendor |
| --- | --- | --- | --- |
| CD3 | Alexa Fluor® 700 | SP34-2 | BD Biosciences |
| CD4 | Brilliant Violet 785™ | OKT4 | Biolegend® |
| CD4 | fluorescein isothiocyanate | OKT4 | Biolegend® |
| CD8 | V500 | RPA-T8 | BD Biosciences |
| CD45 | BD Horizon™ BV450 | D058-1283 | BD Biosciences |
| CD45 | PE | D058-1283 | BD Biosciences |
| CD45RA | PE-Cy™7 | 5H9 | BD Biosciences |
| CCR5 | PE | 3A9 | BD Biosciences |
| CCR7 | PE-CF594 | 150503 | BD Biosciences |
| CCR7 | Alexa Fluor® 647 | 150503 | BD Biosciences |
| CD69 | Brilliant Violet 650™ | FN50 | Biolegend® |
| CD69 | PE-Cy™7 | FN50 | BD Biosciences |
| CD103 | APC | Ber-ACT8 | Biolegend® |
| IFNγ | APC | B27 | BD Biosciences |
| IL-2 | PE-Cy™7 | MQ1-17H12 | BD Biosciences |
| TNFα | fluorescein isothiocyanate | Mab11 | BD Biosciences |

### Mouse antibodies

Mouse antibodies:
| antibody | conjugate | clone | vendor |
| --- | --- | --- | --- |
| CD3e | PE-CF594 | 145-2C11 | BD Biosciences |
| CD4 | Brilliant Violet 711™ | RM4-5 | Biolegend® |
| CD8 | Brilliant Violet 421™ | 53-6.7 | Biolegend® |
| CD44 | Alexa Fluor® 700 | IM7 | Biolegend® |
| CD45.1 | Brilliant Violet 510™ | A20 | Biolegend® |
| CD45.2 | APC | 104 | Biolegend® |
| CD62L | Brilliant Violet 605™ | MEL-14 | Biolegend® |
| CD69 | Alexa Fluor® 488 | H1.2F3 | Biolegend® |
| CD90.2 | Brilliant Violet 570™ | 30-H12 | Biolegend® |
| CD103 | PerCp/Cyanine5.5 | 2E7 | Biolegend® |
| Ly6G | Brilliant Violet 510™ | 1A8 | Biolegend® |
| CX3CR1 | Brilliant Violet 650™ | SA011F11 | Biolegend® |

## Acknowledgements

We thank the study participants and colleagues who contributed to the HC and MVC clinical studies. From Emory University we thank Dr. Bill Schafer for assistance with growing *C. muridarum* cultures and Dr. Rama Amara for providing SHIV_162P3_ envelope peptides. From CDC we thank Dr. Janet McNicholl for assistance in reviewing the manuscript, Dr. Jim Smith, Sunita Sharma, Susan Rhone, Mian-er Cong, Dr. Charles Dobard & Kenji Nishiura for assistance with NHP studies, Cheng-Yen Chen and Kathryn A. Lupoli for STI testing in human samples, James Mitchell, Shanon Ellis, Frank Deyounks, Kristen Kelley for animal technical assistance, and Dr. David Garber for programmatic support. This study was supported by the U.S. Centers for Disease Control and Prevention, Atlanta, GA 30329 and Emory University and in part by NIH grants R35HL150803 (J.E.K) and U01HL139483 (R.A. and J.E.K.), K23AI114407 (A.N.S), K23HD078153-01A1 (L.B.H), the Emory University Center for AIDS research (P30AI050409), the Atlanta Clinical and Translational Sciences Institute (KLR2TR000455, UL1TR000454), and the Biomarker Core Laboratory at the Yerkes National Primate Research Center Base Grant P51OD011132. This study was supported in part by the Emory Multiplexed Immunoassay Core (EMIC), which is subsidized by the Emory University School of Medicine and is one of the Emory Integrated Core Facilities. Additional support was provided by the National Center for Georgia Clinical & Translational Science Alliance of the National Institutes of Health under Award Number UL1TR002378.

## Author Contributions

A.S-K. supervised, designed and performed experiments, analyzed data, and wrote the paper with assistance from all authors. A.N.W. performed mouse microscopy experiments. A.N.S. and L.B.H. designed human studies. E.M.B.C. supervised interpretations from human biopsies and hysterectomies. M.E.W. performed bioinformatic analysis, J.R-B. and H.Z. assisted with NHP experimentation. F.B.H. and Z.T.L. performed statistical analysis and mathematical modeling. Z.T.L. optimized ddPCR approach. C.D. performed MVC drug measurements. S.H., K.K, J.L.L, and Z.T.L., assisted with mouse experiments. A.C.L. supervised virus engineering. M.E.W. performed bioinformatic analysis, C.D.S. supervised genomic studies. I.O. supervised the human study. R.A. supervised mathematical modeling. G.G-L. Supervised and designed the NHP studies. J.E.K. supervised, designed experiments, analyzed data, and wrote the paper with assistance from all authors.

## Competing interests

The authors declare no conflict of interest.

## Disclaimer

The findings and conclusions in this report are those of the authors and do not necessarily represent the views of the U.S. Centers for Disease Control and Prevention, the Department of Health and Human Services, or the National Institutes of Health.

## Notes

### Competing Interest Statement

The authors have declared no competing interest.

