## Supplemental data for "Memory CD4 T cell subset organization in the female reproductive tract is regulated via the menstrual cycle through CCR5 signaling"

**Figure S1. Chemokine receptor expression from FRT CCR7$^{hi/lo}$ memory CD4 T cells compared with paired PBMC.** Indicated chemokine receptors or integrins was measured from $T_{MM}$: CCR7$^{hi}$ CD45RA$^{lo}$ or $T_{RM}$: CCR7$^{lo}$ CD45RA$^{lo}$ gated populations of FRT (open circles) or PBMC (black circles) CD4 T cells (N≥9 samples tested for each marker group). *p<0.05, **p<0.01, ***p<0.001, ****p<0.0001.

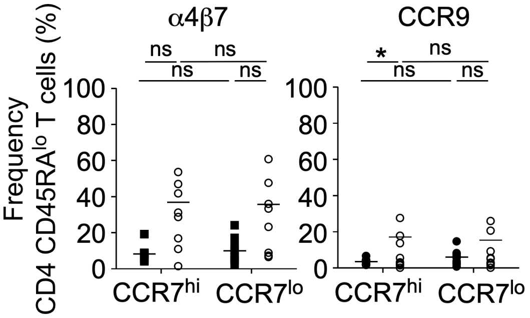

**Figure S2. (A).** Profile of the female human form created with BioRender.com. (B). Representative gating strategy for intracellular measurement of IFNγ, IL-2, TNFα, and IL-17A from activated $T_{MM}$: $CCR7^{hi}$ $CD45RA^{lo}$ (top) or $T_{RM}$: $CCR7^{lo}$ $CD45RA^{lo}$ (bottom) CD4 T cell populations enriched from CVL. **(C).** The frequency of IFNγ, IL-2, TNFα, and IL-17A production measured from $CCR7^{hi}$ $CD45RA^{lo}$ or $CCR7^{lo}$ $CD45RA^{lo}$ CVL enriched CD4 T cells (N=9, graph depicted as a column plot with error bar**).** Means evaluated by multiple comparisons. **(C).** *p≤0.05, **p<0.01, ***p<0.001, ****p<0.0001.

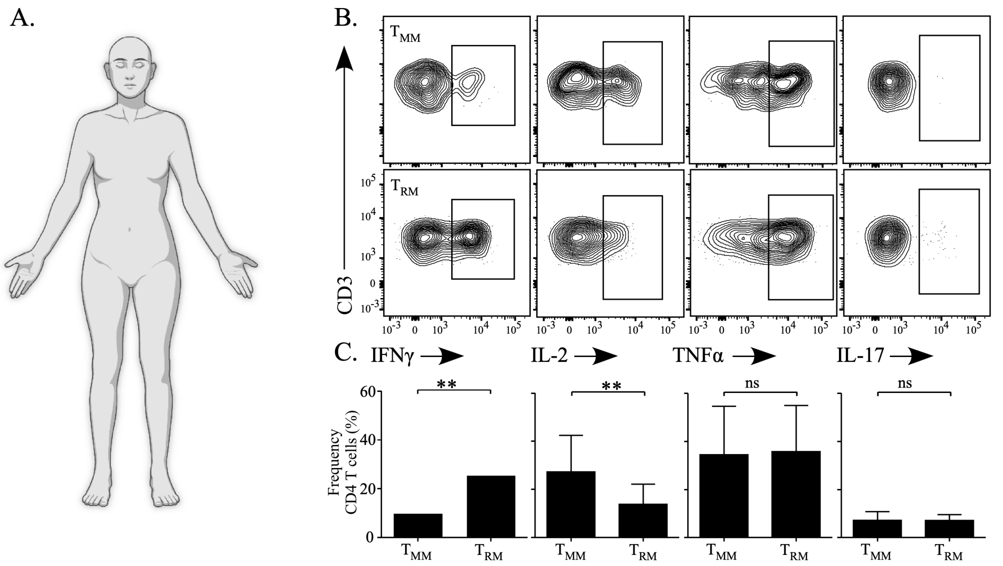

**Figure S3. (A).** Cartoon of a rhesus macaque created with BioRender.com**. (B).** Weekly measurements of plasma $SHIV_{162P3}$ RNA from the time point of seroconversion in female rhesus macaques (N=3) following rectal challenge. **(C).** The frequency of $CCR7^{hi}$ (white circles) or $CCR7^{lo}$ (black circles) expression of $CD45RA^{lo}$ CD4 T cells from PBMC, CVL, and FRT tissues of $SHIV_{162P3}$ infected macaques (graph depicted as a scatter plot with mean line). **(D).** Pie chart indicating the proportion of IFNγ, IL-2, and TNFα cytokine production from either PBMC, CVL, or FRT-enriched CD4 T cells following $SHIV_{162P3}$ peptide stimulation. **(C, D).** Means evaluated by multiple comparisons. *p≤0.05, **p<0.01, ***p<0.001, ****p<0.0001.

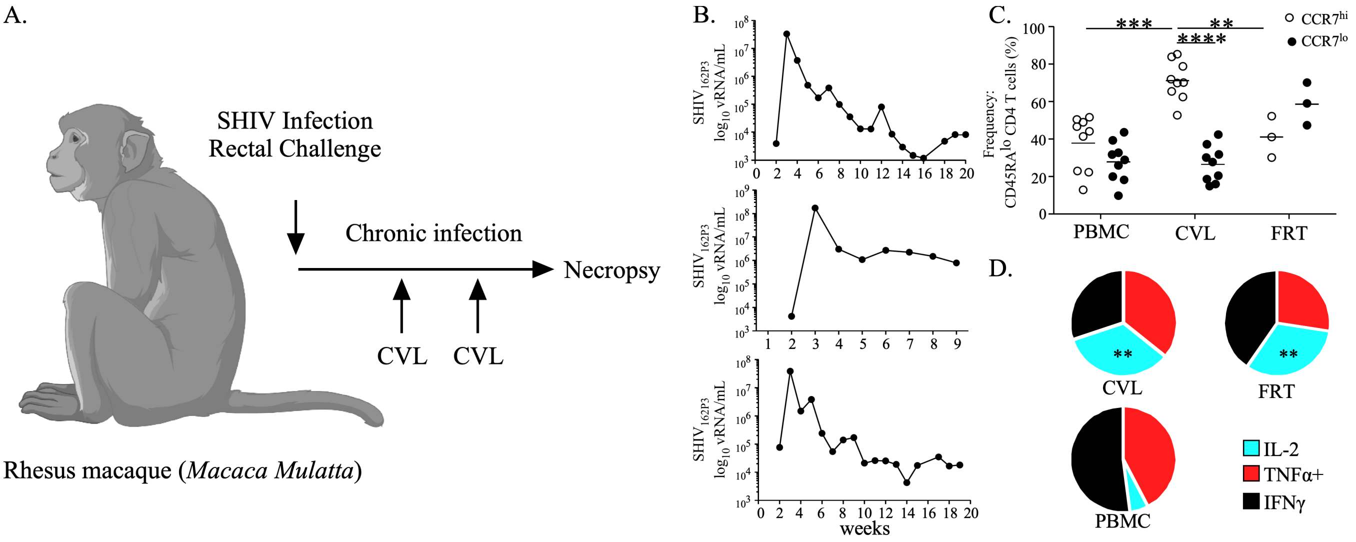

**Figure S4. CD4 T cell intracellular cytokine expression following incubation with overlapping peptide to $SHIV_{162P3}$.** The frequency of IFNγ, IL-2, and TNFα intracellular cytokine expression measured from CVL or FRT-enriched CD4 T cells compared with matched cells from PBMC. Significance calculated by an unpaired t-test *p<0.05, **p<0.01, ***p<0.001, ****p<0.0001 (N=3).

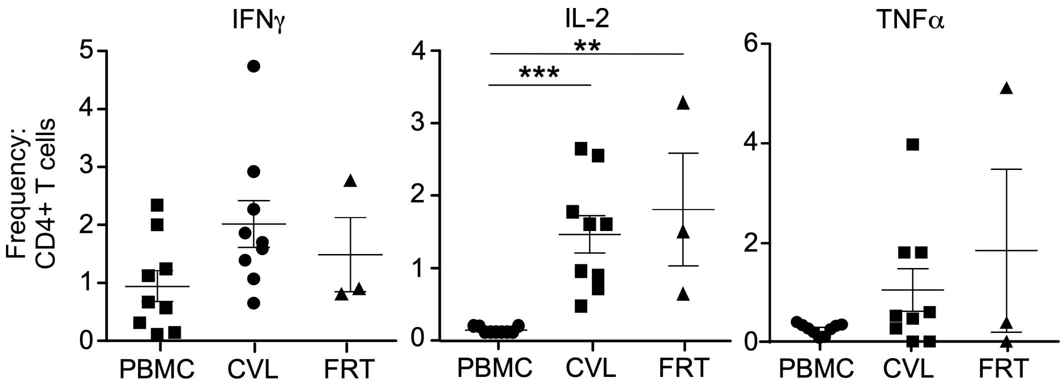

### Table S1. Gene Ontogeny Analysis

| | *Homo sapiens - REFLIST (20595)* | *fold Enrichment* | *raw P-value* | *NEG Log10 P-value* | *False Discovery Rate (FDR)* |
|---|---|---|---|---|---|
| *Cluster I* | hematopoietic or lymphoid organ development (GO:0048534) | 3.8 | 3.07E-09 | 8.51E+00 | 9.75E-06 |
| | regulation of cytokine production (GO:0001817) | 2.8 | 8.26E-06 | 5.08E+00 | 8.19E-03 |
| | cell surface receptor signaling pathway (GO:0007166) | 1.79 | 3.89E-05 | 4.41E+00 | 2.37E-02 |
| | regulation of cell migration (GO:0030334) | 2.41 | 7.92E-05 | 4.10E+00 | 3.22E-02 |
| | regulation of cellular extravasation (GO:0002691) | 12.31 | 9.03E-05 | 4.04E+00 | 3.41E-02 |
| *Cluster II* | cytokine-mediated signaling pathway (GO:0019221) | 4.28 | 2.56E-17 | 1.66E+01 | 4.07E-13 |
| | positive regulation of cell migration (GO:0030335) | 3.67 | 7.87E-10 | 9.10E+00 | 3.57E-07 |
| | positive regulation of T cell activation (GO:0050870) | 4.73 | 3.28E-07 | 6.48E+00 | 5.85E-05 |
| | positive regulation of interferon-gamma production (GO:0032729) | 7.95 | 4.64E-06 | 5.33E+00 | 6.34E-04 |
| | regulation of T cell differentiation (GO:0045580) | 4.9 | 1.27E-05 | 4.90E+00 | 1.47E-03 |
| | cellular response to tumor necrosis factor (GO:0071356) | 3.39 | 1.16E-04 | 3.94E+00 | 8.53E-03 |
| | positive regulation of alpha-beta T cell activation (GO:0046635) | 6.89 | 1.23E-04 | 3.91E+00 | 9.00E-03 |
| | positive regulation of T-helper 1 type immune response (GO:0002827) | 17.16 | 1.81E-04 | 3.74E+00 | 1.19E-02 |
| | positive regulation of CD4-positive, alpha-beta T cell activation (GO:2000516) | 8.83 | 4.18E-04 | 3.38E+00 | 2.38E-02 |
| | positive regulation of T cell proliferation (GO:0042102) | 4.71 | 4.60E-04 | 3.34E+00 | 2.58E-02 |
| | regulation of interleukin-2 production (GO:0032663) | 5.91 | 7.97E-04 | 3.10E+00 | 3.85E-02 |
| | T cell differentiation (GO:0030217) | 3.73 | 7.97E-04 | 3.00E+00 | 4.63E-02 |

**Figure S5. CD4 T cell CCR7-mediated chemotaxis from indicated regions of the FGT**. Mice were intravenously labeled with anti-CD3ε prior to tissue harvest and FRT was dissected into the lower, cervical, and upper region for analysis. CD3ε IV negative CD4 T cells enriched from indicated FGT compartments were measured for CCL21 chemotaxis via trans-well. The chemotactic index was calculated according to the absolute number of CD4 T cells detected in wells incubated with CCL21 and normalized to those in media alone. Graph represent 2 of 2 separate experiments (N=10).

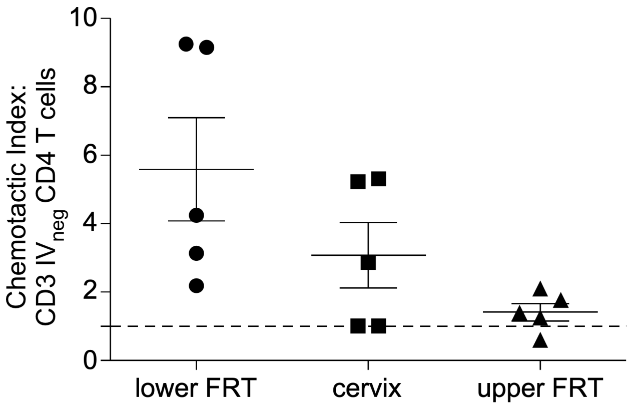

**Table S2.** Cytokine concentrations compared by phase of the menstrual cycle. Statistics calculated by a generalized estimating equation (GEE).

| ***Cytokine*** | ***Comparison*** | ***Mean difference*** | ***Upper 95%*** | ***Lower 95%*** | ***P-value*** |
|---|---|---|---|---|---|
| *IL-8* | Luteal with Follicular | 10417.71 | 5550.24 | 15285.19 | <0.0001 |
| *Mip1a* | Luteal with Follicular | 3.21 | -0.15 | 6.57 | 0.0612 |
| *Mip1B* | Luteal with Follicular | 13.12 | 5.45 | 20.78 | 0.0008 |

**Figure S6. CCR5 regulates steady-state CD4 T cell trafficking into the FRT barrier following $T_{MM}$ priming**.

**(A).** Experimental mouse model schematic depicting the mixed bone marrow chimera approach. Bone marrow cells from CCR5 knockout (KO) and wild type (wt) were transferred at a 1:1 ratio intravenously (IV) into irradiated host mice. Once recovered, chimeras were immunized using a prime boost strategy to induce PmpG-1-specific CD4 T cells. Prior to tissue harvest mice were injected intravenously with anti-CD3ε to discern T cells in the circulation versus tissues. **(B).** The ratio of wt and CCR5 KO CD3ε IV negative CD4 T cells (black circles) or tetramer positive CD4 T cells (white circles) from the indicated tissues (N=10). Models used to evaluate multiple comparisons were fit using a two-way ANOVA, p-values ≤0.05 are shown as *p≤0.05, **p<0.01, ***p<0.001, ****p<0.0001. Graphs depicted as scatter plot with mean line.

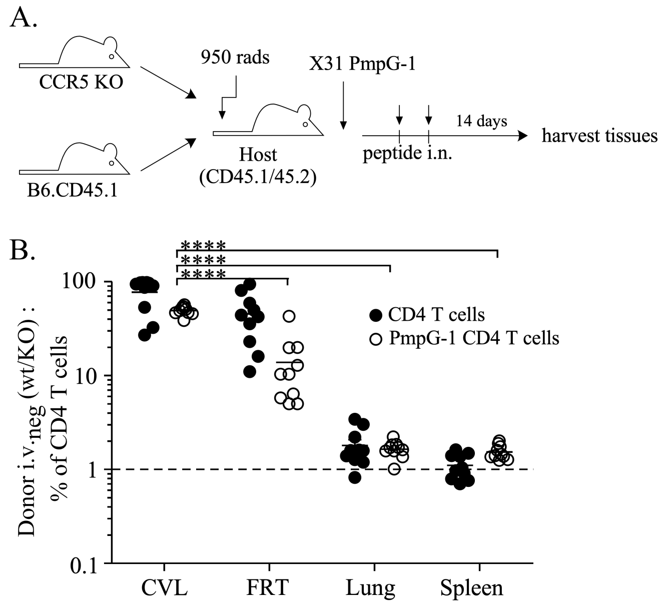

**Figure S7. CD4 T cells in the FGT of pigtail macaques following MVC treatment.** (Left panel) CVL enriched CD4 T cells were measured for the estimated absolute number of cells (determined by BD Trucount™ estimation) during placebo time points during the relative follicular or luteal phase of the menstrual cycle. (Center panel) CVL enriched CD4 T cells measured for the estimated absolute number of cells during placebo time points at Day 5 compared to Day 0. (Right panel) CVL enriched CD4 T cells was measured for the estimated absolute number of cells during Maraviroc treatment at Day 5 of treatment compared to Day 0. Significance calculated by unpaired t-test (all comparisons were found not significant; P>0.05).

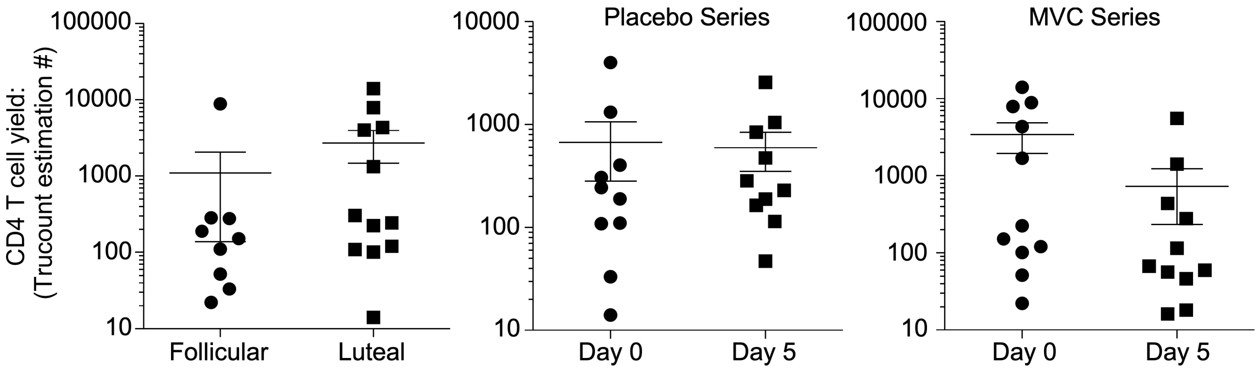
